# Rhythmic Nuclear Import Mediated by Importins Regulates the *Neurospora* Circadian Clock

**DOI:** 10.64898/2026.01.12.699091

**Authors:** Ziyan Wang, Bin Wang, Bradley M. Bartholomai, Jennifer J. Loros, Jay C. Dunlap

## Abstract

Circadian clocks in eukaryotes rely on precisely regulated negative feedback loops to generate daily rhythms. However, the delay mechanisms that extend this structurally simple feedback loop to ∼24 hours are not yet fully understood. In the filamentous fungal model organism *Neurospora crassa*, the negative arm complex, centered by FREQUENCY (FRQ), must enter the nucleus to repress the White Collar Complex (WCC) and close the feedback loop, but the mechanisms and dynamics of its nuclear transport have remained unresolved. Using long-term live-cell imaging and fluorescence recovery after photobleaching (FRAP), we demonstrate that FRQ nuclear import is an active circadian-regulated process that is fastest early in the subjective day and whose rate progressively decreases as nuclear FRQ approaches peak levels, corresponding to altered direct binding between FRQ and Importin α. We further establish that Importin α is required for the spatial regulation of FRQ and WCC, as well as the correct timing of *Neurospora* circadian clock, whereas the nuclear accumulation of non-clock-related free WC-2 doesn’t require Importin α. Analysis of the three *Neurospora* Importin β homologs reveals that each of them contributes differently to the circadian clock through pathways beyond FRQ or WCC nuclear import. More specifically, we find a genetic interaction between Impβ3 and the phosphatase PPH-4. Together, these findings indicate that nuclear import is a selective, dynamic, and rate-limiting regulatory step in the fungal circadian clock and reveal both conserved and fungal-specific mechanisms by which importins tune circadian timing.

## INTRODUCTION

Circadian clocks are self-sustained timekeeping systems that allow organisms to anticipate and adapt to daily changes in the environment. These oscillators are based on conserved transcription-translation feedback loops (TTFLs) in which the Positive Arm (typically transcription factors) drives expression of the Negative Arm that, once translated and imported into nuclei, inhibit its own activators (Dunlap, 1999; Hardin, 2011; Partch et al., 2014). A key feature of this design is that multiple built-in delays extend the simple feedback loop to maintain a ∼24-hour periodicity that approximates environmental changes.

Precisely timed nuclear entry of Negative Arm components can act as a rate-limiting step that introduces phase delay between transcriptional activation and repression. Importins (karyopherins) mediate the nucleocytoplasmic transport of most proteins through the nuclear pore complex (NPC) (Chook and Blobel 2001; Pemberton and Paschal 2005). There are two major nuclear import pathways: the classical Importin α (Impα)-dependent pathway, in which Impα, as an adaptor, binds cargos with canonical nuclear localization signals (NLSs) and delivers them to Importin β (Impβ); and the Impα-independent pathway, in which Impβ directly recognizes non-classical NLSs. In *Drosophila*, the clock protein TIMELESS binds directly to Impα1 and co-transports Drosophila PERIOD (dPER), and perturbations of Impα1 alter circadian period and rhythmicity (Jang et al. 2015; Lin et al. 2023). In mammals, the extent to which nuclear import of PERs and CRYs requires Impα remains debatable and may rely more directly on Impβ family members. In human cells, KPNB1 (Impβ1) mediates nuclear import of some PER/CRY complexes independently of Impα (KPNA1/2), and TNPO1 (Impβ2) selectively regulates PER1 localization and period length (Lee et al. 2015; Korge et al. 2018; Lee et al. 2019). However, chemical inhibition of the classical Impα/β1 pathway also alters period length (Öllinger et al. 2014), and in mouse, nuclear import of mCRY2 is Impα-dependent (Sakakida et al. 2005). These findings suggest that controlled nuclear entry of the Negative Arm is a conserved step important for circadian timing, but the exact pathways and molecular players differ: Impα is essential in *Drosophila*, while in mammals PER and CRY can sometimes bypass Impα and depend on specific Impβs. In contrast to animal circadian systems, little is known about how the Positive and Negative Arm components of fungal clocks such as *Neurospora* enter the nucleus.

The filamentous fungus *Neurospora crassa* has been a powerful model for studying circadian clocks. The Positive Arm of its oscillator is the WCC, a heterodimer of WC-1 and WC-2, which activates transcription of the *frequency* (*frq* gene) whose product serves to nucleate the Negative Arm complex of FRQ, FRQ-interacting RNA helicase (FRH), and Casein Kinase 1(CK1). This complex then feeds back to repress WCC activity (Crosthwaite et al. 1997; Dunlap 1999; Dunlap and Loros 2017; Costa et al. 2025). FRQ is highly concentrated in the nucleus, which makes sense because its nuclear entry is required for inhibitory function on WCC and is essential for closing the negative feedback loop (Wang et al. 2026). Thus, FRQ nuclear transport is a central regulatory point in the *Neurospora* clock.

Despite its importance, the mechanism of FRQ nuclear transport remains unclear. An NLS (aa 194-199, PRRKKR) within FRQ was reported more than two decades ago and suggested to be essential for both nuclear localization and clock function (Luo et al. 1998), but this has not been thoroughly tested with modern tools, and the transport machinery has also not been defined. Unlike animals, which express multiple Importin α isoforms (e.g. seven paralogs, KPNA1-7, encoded by a multigene family in humans) with overlapping and specialized functions, plants and fungi only have one class of Importin α that most closely resembles animal α1 (Goldfarb et al. 2004; Mason et al. 2009; Bernardes et al. 2020). Within the fungal kingdom, nuclear import pathways have only been systematically studied in yeast and *Aspergillus nidulans*. A few related studies in *Neurospora* focused on the conserved structure and epigenetic function of Impα (also referred to as DIM-3, encoded by *dim-3*/NCU01249) (Takeda et al. 2013; Bernardes et al. 2015; Klocko et al. 2015). Through identification of homologs, using NCBI orthologs based on the Karyopherin-β family in *Saccharomyces cerevisiae* and *Aspergillus nidulans* (Ström and Weis 2001; Mans et al. 2004; Markina-Iñarrairaegui et al. 2011), we identified a single Impα, three Impβs and nine other potential Karyopherins including Exportin-1 in *Neurospora crassa* (Supplementary Table 1). Mass spectrometry studies suggested FRQ may weakly interact with Impα and/or Impβ3 (Baker 2010; Pelham et al. 2023), but biochemical, functional and biological validation is lacking. How these Importins contribute to FRQ nuclear entry or to subcellular localization of other clock proteins is unknown. In part this is due to technical challenges which have further limited progress. The insufficient brightness of FRQ-GFP fusion protein prevented reliable imaging, and an “anchor-away” assay (Haruki et al. 2008), which used FRQ fused to FKBP-rapamycin-binding domain of mTOR (FRB) artificially tethered to FK506-binding protein (FKBP)-tagged histone H1 to assay nuclear import, introduced non-physiological import routes and disrupted normal clock function (Diernfellner et al., 2009). As a result, the dynamics and regulation of FRQ nuclear transport in *Neurospora* remain poorly characterized.

In this study, we used live-cell imaging combined with FRAP to directly measure FRQ nuclear import in *Neurospora*. We found that FRQ is imported slowly, requiring tens of minutes for recovery, and that import is strongly regulated across the circadian cycle, being highest early in the subjective day and reduced near nuclear FRQ peak levels. These dynamics are comparable to those of mammalian PER proteins, suggesting that slow nuclear import is a conserved delay mechanism in circadian clocks. We further show that FRQ import requires Impα but not the previously proposed NLS, and that import of WC-1, but not WC-2, also depends on Importin α. Finally, we find that the three importin βs have distinct contributions to circadian regulation, including a potential link between Impβ3 and PPH-4. Together, these results reveal nuclear import as a selective and regulated process that is essential for circadian timing in *Neurospora*.

## RESULTS

### Nuclear import rate of FRQ is circadianly regulated

Precisely timed nuclear localization of FRQ is essential for the normal operation of the circadian clock. To understand how and when FRQ enters nuclei, we optimized FRAP experiments for *Neurospora* building on the continuous live-cell imaging approach reported separately (Wang et al. 2026). Briefly, a slow-growing, light-blind strain carrying fluorescent tags and an intact circadian clock can be cultured in customized microfluidic devices for multi-day imaging of circadian dynamics after temperature resetting of the clock (28°C and 21°C representing day/light-on and night/light-off, respectively). In this study, we used an imaging strain blind to blue light carrying endogenously mNeonGreen (mNG)-tagged FRQ (strain 1952-2; *his-3::frq_cbox_-luc; ras-1^bd^; wc-1*^Δ*LOV*^*, for, frq^mNeonGreen_Flag^*); this strain retains an intact circadian clock (Fig. S1a-c), with the tagging strategy validated in the previous publication (Wang et al. 2026). The application of microfluidic devices allows convenient medium exchange, enabling application of the microtubule inhibitor benomyl at specific times. Benomyl slows nuclear movement by disrupting active transport along microtubules and by reducing growth, which decreases passive movement driven by bulk cytoplasmic flow and thereby facilitates imaging. Importantly, benomyl does not affect circadian functions (Fig. S1d). For FRAP experiments, a thick optical section (around 2 μm, corresponding to the diameter of *Neurospora* nuclei) was chosen to minimize artifacts from vertical random movement of nuclei. Cultures in microfluidic devices (Wang et al. 2026) were entrained with 12h:12h temperature cycles mimicking natural conditions, and FRAP was then performed every 30 minutes during the subjective day on the same culture (Fig.1a). After bleaching entire target nuclei, the subsequent recovery of fluorescence reflects import of unbleached cytoplasmic FRQ^mNG^ at that time.

The nuclear fluorescence from FRQ^mNeonGreen^ recovered most rapidly early in the subjective day, indicating a higher import rate, while the recovery was slower later in the subjective day (Fig. 1b-c, Supplementary Video 1). This temporal difference is consistent with the progressive nuclear accumulation of FRQ during the subjective day that we observed previously (Wang et al. 2026). Near subjective dawn, the recovered signal could even exceed the pre-bleaching level (e.g. 129 ± 29% at 23 minutes post-bleaching at CT3; CT, circadian time). By contrast, at subjective dusk, around when nuclear FRQ levels reach their peak, nuclear import was remarkably reduced (Fig. 1c-d).

**Figure 1.**
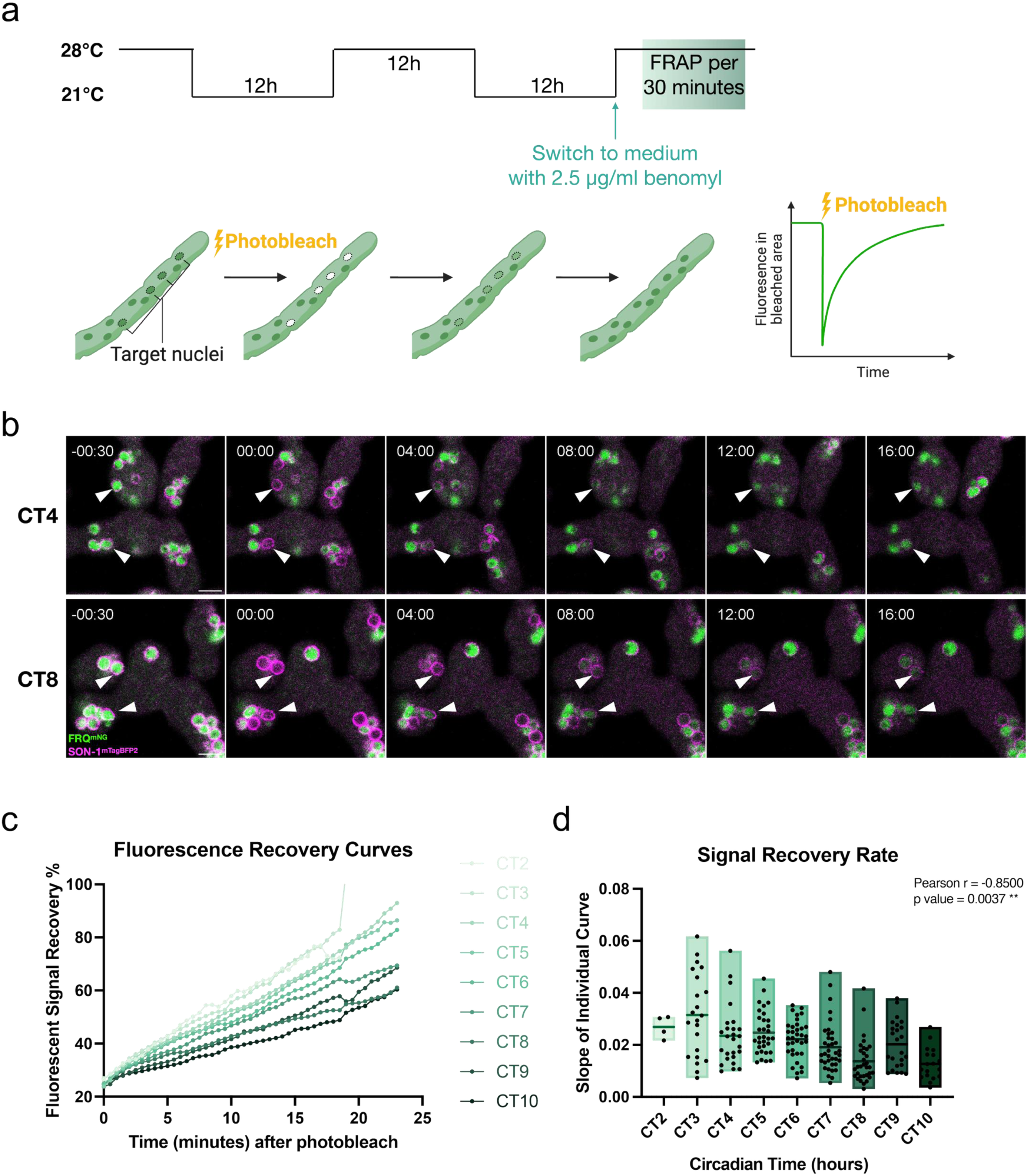
Nuclear import of FRQ is circadian regulated. a) Schematic of the experimental design. (Top) FRAP experiments were performed during the subjective day following entrainment by temperature cycles. (Bottom) Whole nuclei were bleached, and the recovery of their fluorescent signal reflects nuclear import of cytoplasmic FRQ. b) Example timelapses from subjective morning (CT4, top) and afternoon (CT8, bottom), showing different recovery rates. CT: circadian time. White arrowheads indicate example target nuclei. Scale bar = 5 μm. c) Average fluorescent recovery curves for each hour of the subjective day. The variations (standard deviations) for each curve are included in Fig. S1e. d) Nuclear signal recovery speed, calculated as the slope of linear regression of each target nucleus’s recovery curve. The floating box is showing the minimum to maximum range, with a line at the mean. Data were analyzed by Pearson correlation, with the correlation coefficient (r) and p-value indicated in the plot.

Due to the slow FRQ import rate we observed, most recovery curves failed to reach a stable plateau within the 25-minute observation window. To ensure consistent analyses, we used linear regression rather than the commonly used exponential fit to calculate nuclear import rates. To compensate for high variability in nuclear movements and possible artifacts, multiple control nuclei within each culture were averaged and used to correct for unavoidable photobleaching caused by continuously imaging. The nuclear import rate shows a significant negative correlation with time during the subjective day (Fig. 1d, Fig. S1e-f). Consistent results were observed when using a single corresponding nucleus from the same experiment, without averaging (Fig. S1g). In conclusion, nuclear import rates of FRQ are circadian regulated with a peak in the subjective morning.

### FRQ is transported into nuclei through the Impα import pathway

Because previous mass spectrometry had identified Impα as a possible weak interactor of FRQ (Baker et al. 2009; Baker 2010), we hypothesized that the time-dependent regulation of FRQ nuclear import is mediated by its interaction with Impα. To verify this interaction, we endogenously tagged Impα with the V5 epitope tag at its C-terminus. Co-immunoprecipitation with V5 beads successfully enriched Impα^V5^, and FRQ co-precipitated with Impα^V5^at early time points, corresponding to times showing rapid nuclear import (Fig. 2a, Fig, S2a-b). However, at later time points when FRQ is more abundant, binding between FRQ and Impα^V5^ is barely detectable, corresponding to times of slower import. The change in binding strength is consistent with this being the cause for precisely controlled nuclear import. Phosphorylation status of FRQ is different at CT4 and CT8, as expected, with hypophosphorylated FRQ at CT4 showing a stronger binding with Impα, suggesting that the binding change could be related to an alteration of FRQ conformation due to phosphorylation.

**Figure 2.**
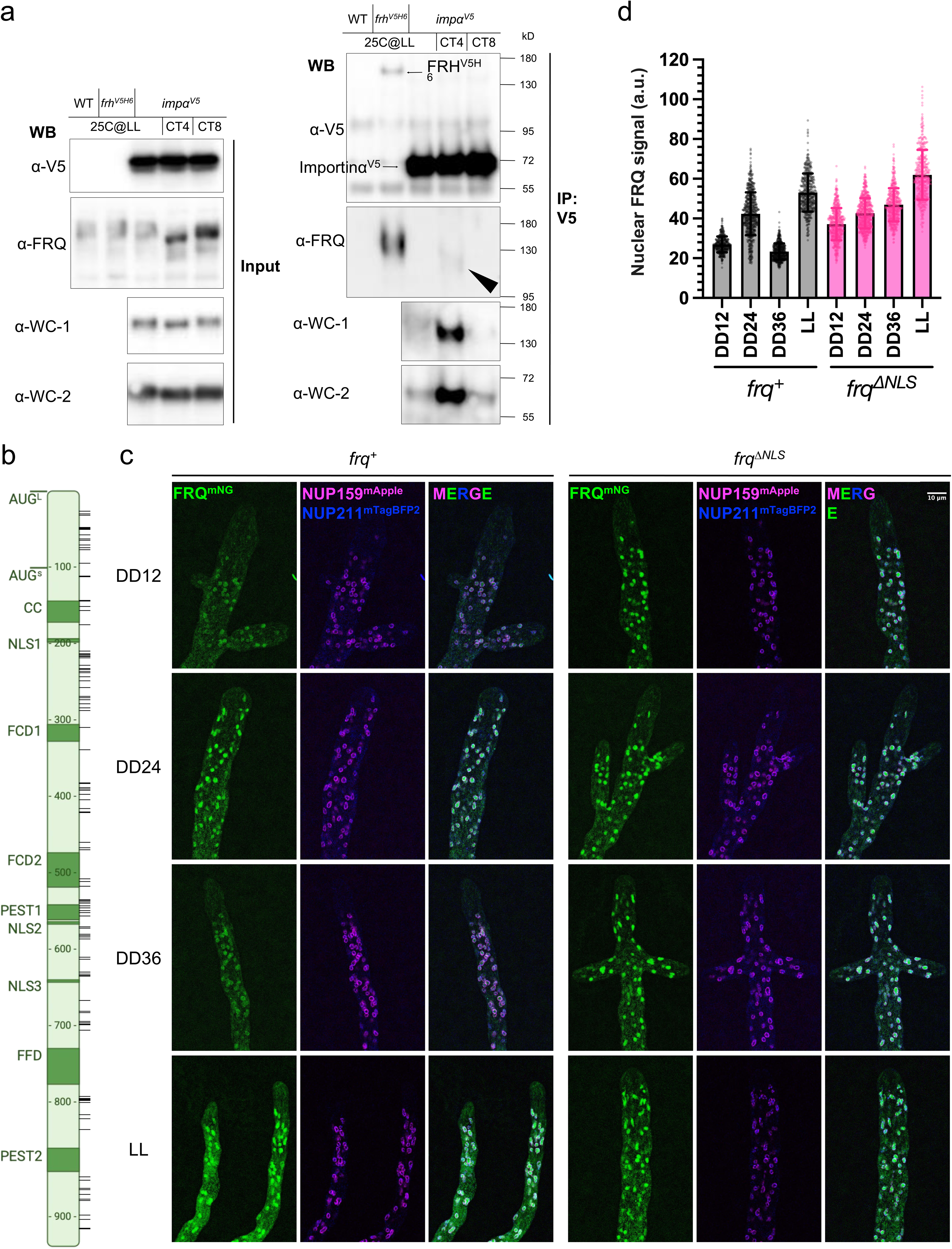
Core clock proteins directly bind Importinα while bypassing the classic NLS for nuclear import. a) Co-immunoprecipitation between endogenous Importin α and core clock proteins. Cell lysates were immunoprecipitated with anti-V5 antibody. The interaction between FRQ and FRH (Lane 2) is included as a positive control for V5-IP based interaction detection; the wildtype strain (Lane 1) is included as a negative control. LL samples were cultured under constant light. b) Schematic of known important domains (dark green) and phosphorylation sites (horizontal black bars) on FRQ. AUG_L_ and AUG_S_, translational start sites of the long and short FRQ isoforms; CC, coiled-coiled domain; NLS#, nuclear localization signal (NLS1, RRKKR; NLS2, RRKKRK; NLS3, RRKRR); FCD#, FRQ-CK1 interacting domains; PEST#, PEST domains. FFD, FRQ-FRH interacting domain. The phosphorylation sites are based on two publications (Baker et al. 2009; Tang et al. 2009). c) Example central focal images showing FRQ or FRQ^ΔNLS^ subcellular localization during free-run circadian cycles or in constant light. FRQ is shown in green, the inner nuclear envelope marker NUP211 in blue, and the outer nuclear envelope marker NUP159 in magenta. All targets are endogenously tagged with monomeric fluorescent proteins. DD time stands for the duration (hours) in constant darkness following a light-to-dark transition. Scale bar = 10 μm. d) Quantification of nuclear FRQ or FRQ^ΔNLS^ signal from 3D renderings of z-stacks (Mean ± SD). Each dot represents the background subtracted mean fluorescent signal of one nucleus. N = 7-11 hyphal tips per condition, with the ungrouped data of each tip plotted in Fig. S2e.

Impα is expected to act through an NLS (Conti et al. 1998; Takeda et al. 2013). Several putative NLSs have been suggested in FRQ, with one of them being essential to the circadian clock (Luo et al. 1998) (Fig. 2b). We reconstituted Neurospora strains bearing FRQ at its endogenous locus with this FRQ-NLS (aa 194-199, PRRKKR) deleted, with or without the mNeonGreen tag to allow for fluorescent imaging, to directly assess the contribution of the predicted NLS to FRQ nuclear entry. The reconstituted strains recapitulate the arrhythmic phenotype as expected (Fig. S2c). To delineate nuclear boundaries, two nuclear envelope proteins, Nup159 (NCU08604) in the cytoplasmic side of nuclear pore complexes and Nup211 (NCU04059) in the nuclear basket complexes, were tagged at their endogenous loci using different fluorescent tags to serve as markers for the outer and inner sides of the nuclear membrane, respectively (Beck and Hurt 2017; Bartholomai 2022). Both proteins form fragmented ring-like structures that largely align with each other at the nuclear periphery, with the two layers clearly distinguishable when imaged on SoRa super-resolution microscope (Fig. S2d). To our surprise, although rhythmicity was plainly diminished (Fig. S2c), FRQ still localized to nuclei and nuclear FRQ levels remained constantly high when the proposed NLS is absent (Fig. 2c-d, Fig, S2e). This nuclear localization is not an artifact caused by mNeonGreen tagging, as other tagged *Neurospora* proteins, like the septin CDC-11, localized to cytoplasm as expected (Fig. S2f). A concurrent study biochemically confirmed the nuclear accumulation of FRQ with this NLS mutated (RRKKR to RQKKQ) using the nuclear fractionation method (Schunke et al. 2026). Nuclear FRQ still forms highly dynamic nuclear bodies in the *frq*^Δ*NLS*^ strain, with subnuclear heterogeneity levels indistinguishable from that of wildtype FRQ (Fig. S2g, Supplementary Video 2).

Impα plays an important role in *Neurospora* development and growth (Klocko et al. 2015) and complete depletion of Impα is lethal. To assess the functional role of Impα, we used an RNAi approach to knock down *imp*α (=*dim-*3) under the control of a quinic acid (QA)-inducible promoter (Giles et al. 1985; Shi et al. 2010) (Fig. 3a). Addition of QA leads to severe growth defects (Fig. 3b), and short induction (20 hours) is sufficient to significantly reduce both *imp*α transcript and Impα protein levels, while not causing macroscopical sickness or death (Fig. 3c-d). Due to leakiness of the *qa-2* promoter (Larrondo et al. 2009), some reduction of Impα was also observed in the inducible RNAi strains even without induction (Fig. 3c-d, Fig, S3a). The progressive decrease in Impα levels, from the no-dsRNA control strain (*imp*α*^V5^*) to inducible RNAi strains without QA, and finally to the inducible RNAi strains with QA, provided a platform to assay dose-dependent circadian phenotypes caused by Impα deficiency.

**Figure 3.**
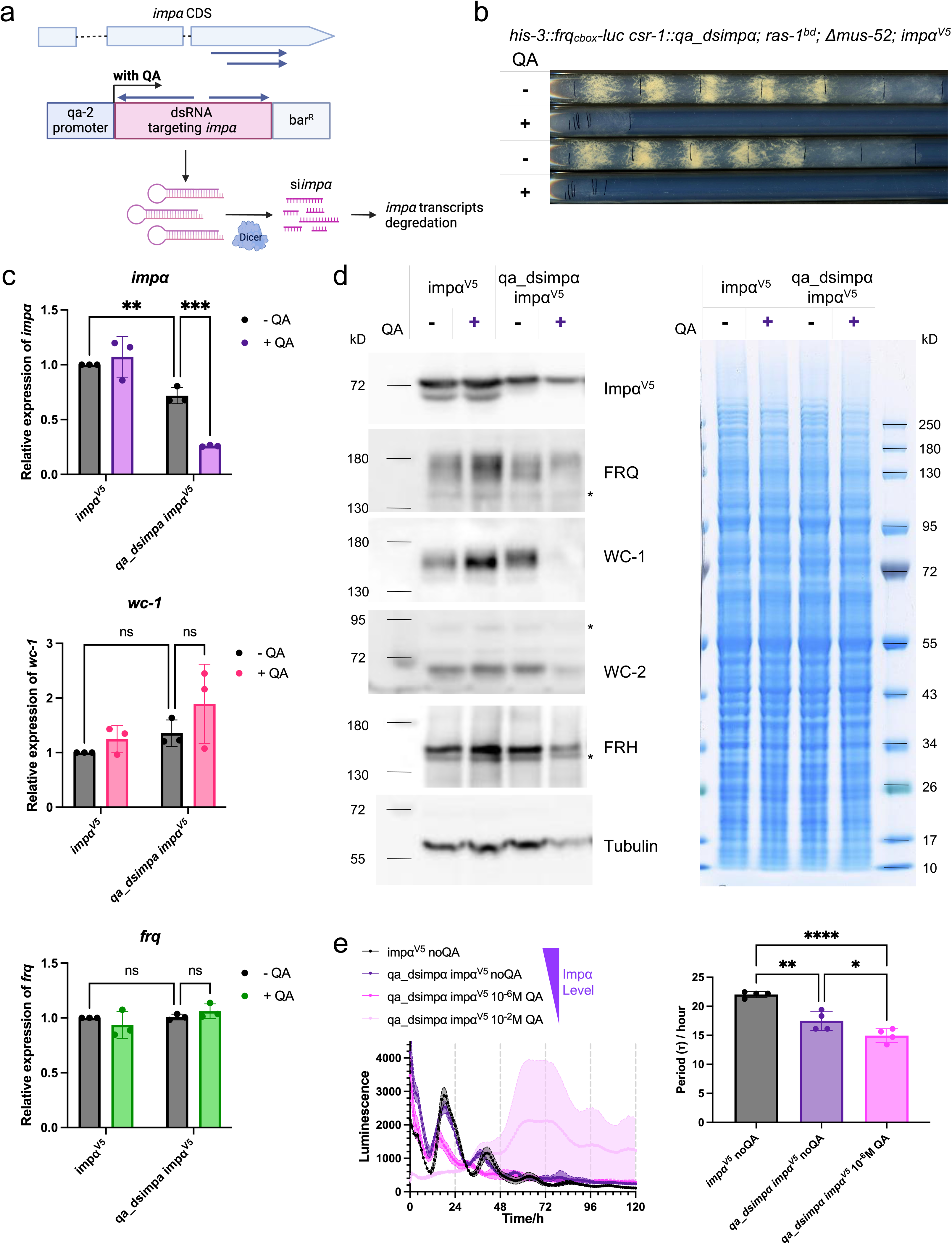
Impα is required for normal circadian rhythmicity. a) Schematic of the siRNA knockdown design. Expression of dsDNA targeting the last exon of *imp*α is driven by the *qa-2* inducible promoter. With the addition of the inducer quinic acid (QA), dsRNA is expressed and subsequently processed to inhibit *imp*α expression. b) Growth of *qa_dsimp*α strains in race tubes with or without 10^-2^ M QA. c) Expression level of *imp*α (top), *wc-1* (middle), or *frq* (bottom) in control (*imp*α*^V5^*) or *qa_dsimp*α strains with or without 10^-2^ M QA quantified by RT-qPCR. Error bars represent the standard deviation from biological triplicates. d) Protein levels in control (*imp*α*^V5^*) or *qa_dsimp*α strains with or without 10^-2^ M QA. Example targets, including core clock proteins, were detected by immunoblotting (left), and total protein levels were assessed by Coomassie staining (right). * indicates unspecific bands in immunoblots. e) Luciferase assays of control (*imp*α*^V5^*) or *qa_dsimp*α strains with varying QA concentrations (top) and corresponding period lengths (bottom). n=4 replicates per condition. Error bars show one standard deviation. Periods were analyzed by one-way ANOVA with multiple comparisons.

We examined circadian rhythms in Impα knockdown strains using the luciferase reporter driven by the *frq_cbox_* promoter described above. Control cultures exhibited ∼22 h rhythms, whereas progressive depletion of Impα shortened the period and stronger knockdown abolished rhythmicity, resulting in arrhythmic or inviable cultures (Fig. 3e, Fig. S3b). Indeed, cultures displayed sensitivity to extremely low levels of inhibition, comparing reductions seen in Fig. 3c-d with rhythm phenotypes in Fig. 3e. Another striking effect caused by Impα deficiency is that all tested core clock proteins, as well as non-clock-related protein Tubulin, decrease in bulk tissues collected from liquid cultures (Fig. 3d). In contrast, the most abundant proteins that dominate unspecific protein measurements remain unchanged. This may reflect a reduction in metabolically active tissue due to sickness in Impα-deficient culture, while the structural components in older tissue remain largely intact.

Live-cell fluorescent imaging allows dissection of local changes in metabolically active tissue, such as the actively growing tip regions. These results are consistent with biochemical analyses, showing that with Impα knocked down, FRQ nuclear import was clearly impaired (Fig. 4a). In the control strains, addition of QA (which also provides an extra nutrient source) boosts nuclear FRQ levels (represented by the mean fluorescent signal in each nucleus) (Fig. 4a-c, Fig. S3c). In contrast, in the RNAi strains, QA addition caused reduced or unchanged nuclear FRQ, compared to the increase in the control strain, despite total FRQ levels in the tip region (hyphal FRQ, represented by the mean fluorescent signal in the whole hyphal tip) remaining relatively constant (Fig. 4a-c, Fig. S3c). This indicates that without sufficient Impα, FRQ accumulates in the cytoplasm and fails to efficiently enter nuclei to shut down its own synthesis. Even in the absence of QA, the leaky knockdown of Impα resulted in a mild reduction in nuclear FRQ compared to controls (Fig. 4b). Together, these observations support the model that FRQ nuclear import is actively mediated by the Impα pathway and is regulated across the circadian cycle to ensure appropriate timing and rates of nuclear entry, both of which are essential for maintaining circadian stability.

**Figure 4.**
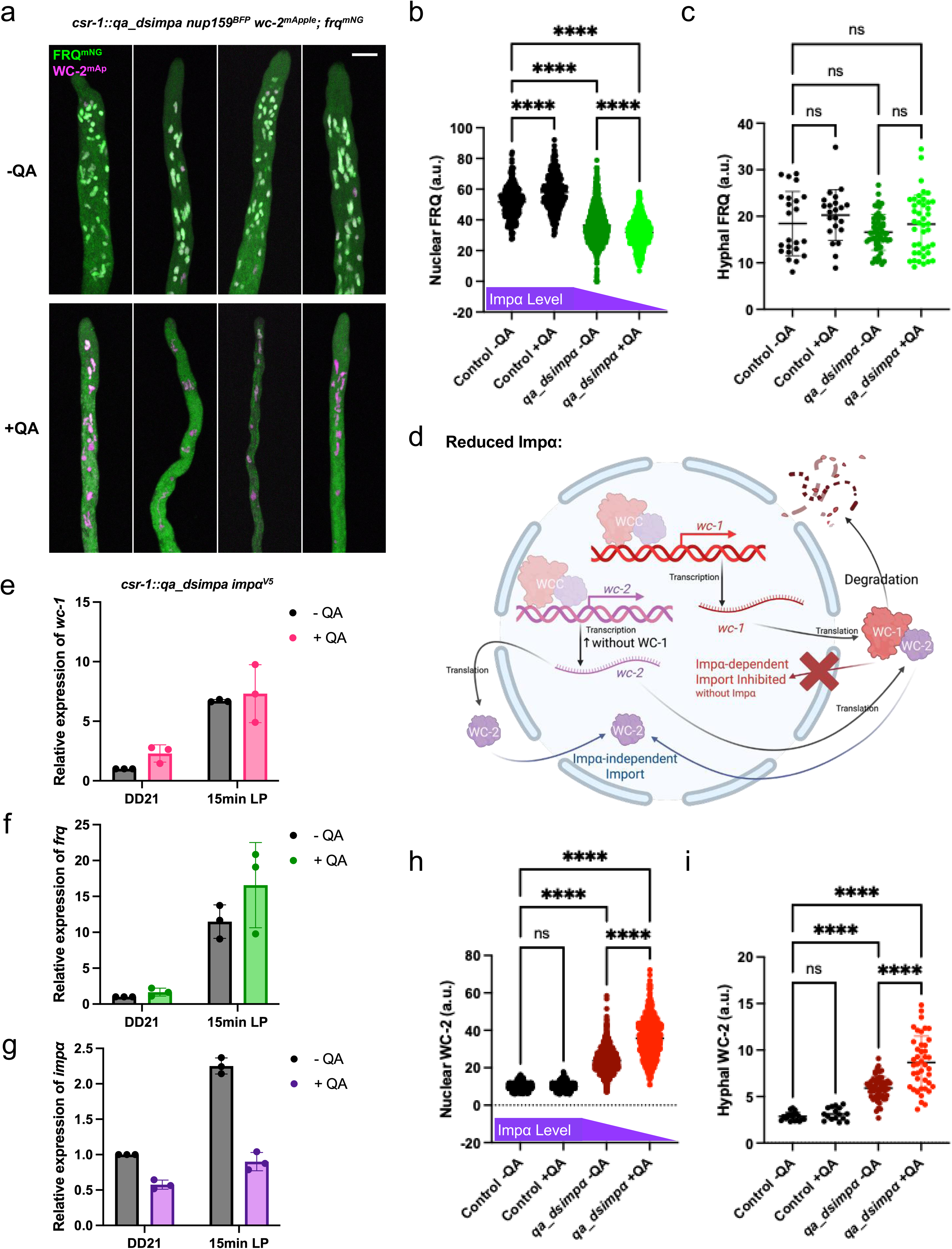
Nuclear localization of FRQ and WC-1, but not WC-2, depends on Impα. a) Maximum projection images of example z-stacks showing FRQ and WC-2 subcellular localization in constant light with or without 10^-2^ M QA. Scale bar = 10 μm. b-c) Quantification of b) nuclear FRQ or C) total FRQ signal in tip regions from 3D renderings of z-stacks. One control strain (1984-4, *nup159^mTagBFP2^; frq^mNG^*) and 2 *csr-1::qa_dsimp*α *nup159^BFP^ wc-2^mApple^; frq^mNG^* strains were analyzed. 2 (controls) or 3 (*dsimp*α strains) biological replicates were quantified, with individual replicates plotted in Fig. S3b. N = 5-12 hyphal tips per condition per replicate. Each dot represents the mean fluorescent signal of one nucleus (nuclear signal) or one hypha (total signal). d) Schematic of the potential mechanism by which WCC amount and localization are affected when Impα is deficient. When WC-1 is trapped in cytoplasm due to lack of Impα, it becomes unstable, while free WC-2, can get imported through other Impα-independent pathway(s). Lack of nuclear WC-1, as a *wc-2* repressor, leads to upregulated *wc-2* expression and thus increased WC-2. e-g) Expression level of e) *wc-1*, f) *frq*, or g) *imp*α in a *qa_dsimp*α strain at DD21 with or without 10^-2^ M QA, before or after 15-minute light pulse, quantified by RT-qPCR. Error bars represent the standard deviation from biological triplicates. h-i) Quantification of h) nuclear WC-2 or i) total WC-2 signal in tip regions from 3D renderings of z-stacks. One control strain (1996-2, *wc-2^mApple^::hph+, his-3::cbox-luc*) and 2 *csr-1::qa_dsimp*α *nup159^BFP^ wc-2^mApple^; frq^mNG^* strains were analyzed. Same methods as Fig. 4b-c were used.

### WCC but not free WC-2 also depends on Impα for nuclear import

The most obvious change in clock protein levels is that WC-1 becomes undetectable by western blot with severe deficiency of Impα (Fig. 3d), even though the expression level of *wc-1* transcripts is unchanged (Fig. 3c). This divergence indicates that the defect occurs at the post-transcriptional level, either through impaired translation or reduced WC-1 protein stability. Interestingly, WC-1 co-immunoprecipitates with Impα in a phase-dependent manner (Fig. 2a), similar to FRQ. Because WC-1 is predominantly found in the nucleus, this suggests a model in which WC-1 is unstable in the cytoplasm. When nuclear import of WC-1 is blocked by deficiency of Impα, newly synthesized WC-1 is trapped in cytoplasm, being unstable, and is rapidly degraded (Fig. 4d). Since WC-1 is always in WCC under normal physiological conditions (Denault et al. 2001; Cheng et al. 2003), the import of WC-1 most likely represents the import of WCC.

Although rhythmic turnover of WCC is not essential for circadian clock regulation, the presence of nuclear WC-1 is essential. While WC-1 is below the detection limit of western blotting in Impα knockdown strains (Fig. S3a), a trace amount of WC-1 may still be able to support circadian function (Lee et al. 2003; Káldi et al. 2006). Consistent with this, the expression of its direct target *frq* is unaffected (Fig. 3c). The photoreceptor function of WC-1 is intact as well; both *wc-1* and *frq* transcript levels increase drastically after a 15-minute light pulse (Fig. 4e-f). Surprisingly, *imp*α itself also turns out to be light-inducible (Fig. 4g), although it was not previously identified as a light responsive gene by RNA-seq (Chen et al. 2009; Smith et al. 2010; Dekhang et al. 2017). Even though the residual WC-1 retains functions when Impα is reduced, its overall efficiency is compromised, as the fold increase of gene expression in response to light is smaller compared to no QA (Fig. S3d). In this context, it may be that less FRQ is needed to fully inhibit the limited WCC activity during circadian cycles, which may explain the observed period shortening despite slow FRQ nuclear import.

WC-2 also co-precipitated with Impα in a phase-dependent manner (Fig. 2a). We endogenously tagged WC-2 with mApple (mAp) (Wang et al. 2026) in the inducible *imp*α strain, and verified that all WC-2 expressed by this allele are proper fusion proteins (Fig. S3a). Surprisingly, both nuclear and total levels of WC-2 in tip regions increase significantly as the level of Impα decreases (Fig. 4a, h-i, Fig. S3e). This Impα-deficient condition represents the case in which WC-1 is not available and WCC is not formed, indicating that the nuclear import of free WC-2 does not appear to rely on the Impα pathway (Fig. 4d). The observed co-immunoprecipitation between WC-2 and Impα reflects the Impα-dependent WCC nuclear import under normal physiological conditions, as its pattern is in full accordance with that of WC-1, while the free WC-2 is not directly associated with Impα.

### Deficiency of importin βs have divergent effects on circadian clock

The genes encoding the three identified Impβ subunits in *Neurospora crassa* are NCU02011, NCU03690, and NCU02357 (Supplementary Table 1) (Galagan et al. 2003). Homologs of two of the three Impβs, Impβ1 (NCU02011; mammalian KPNB1) and Impβ2 (NCU03690; mammalian TNPO1), have been associated with nuclear import of Negative Arm components PER1 and PER2 in mammalian systems (Lee et al. 2015; Korge et al. 2018; Lee et al. 2019). Impβ3 (NCU02357; mammalian IPO5) has not been linked to circadian regulation in any system, but was identified as a potential FRQ interactor at DD24 by mass spectrometry (Pelham et al. 2023). We therefore investigated how deficiency of these three Impβs affects the circadian clock using different strategies.

*imp*β*1* is an essential gene - neither homozygous knockouts nor inducible *imp*β*1* strains (with *imp*β*1* expression driven by either *qa-2* or *tcu-1* promoters) are viable. Moreover, RNAi targeting of two distinct regions of the *imp*β*1* gene, using the same strategy applied to *imp*α, failed to cause a strong growth defect and at best only reduced the expression level by around 40% (Fig. S4a). As an alternative approach to achieve deficiency of Impβ1, we tested two Impβ1 inhibitors, Importazole and Ivermectin (Soderholm et al. 2011; Wagstaff et al. 2011). Both drugs cause growth defects, with Importazole having less severe effects in a dose-dependent manner (Fig. S4b). Based on the growth tests, 40μM drugs were used for fluorescent imaging experiments. Importazole inhibits FRQ nuclear localization (Fig. 5a-c), suggesting a potential role of Impβ1 in nuclear import, possibly together with Impα. Ivermectin treatment instead leads to an overall increase in hyphal fluorescence without changing in nuclear signal (Fig. S4c-e), which could result from cytoplasmic accumulation of FRQ or increased autofluorescence due to growth-associated stress.

**Figure 5.**
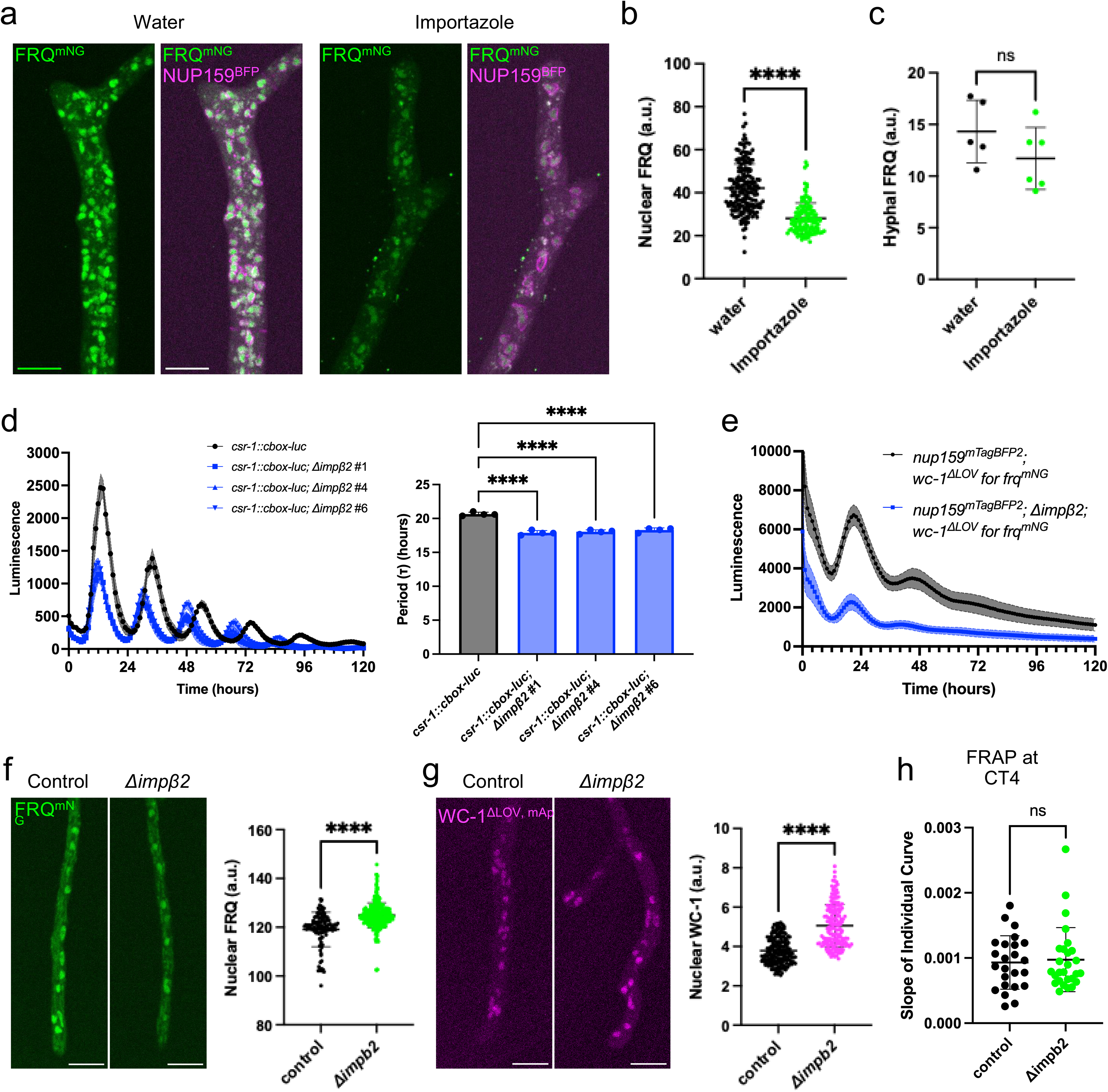
Deficiency of Importinβs affects FRQ localization or circadian clock. a) Maximum projection images of example z-stacks showing FRQ subcellular localization with or without Importazole, an Importinβ1 inhibitor. Samples were treated with drug (or water) and light for 2 hours starting at DD12. N = 5-6 hyphal tips. Scale bar = 10 μm. b-c) Quantification of b) nuclear FRQ or c) total FRQ signal in tip regions from 3D renderings of z-stacks (Mean ± SD). Each dot represents the mean fluorescent signal of one nucleus (nuclear signal) or one hypha (total signal). d) Luciferase assays of control strain (*csr-1::frq_cbox_-luc*) or three Δ*imp*β*2* strains (left) and their period lengths (right). n=4 replicates. The shade shows one standard deviation. Periods were analyzed by one-way ANOVA with multiple comparisons. e) Luciferase assays showing the free-run clock of Δ*imp*β*2* FRQ imaging strain used in Fig. 5h compared to its control strain (2066-4) without Δ*imp*β*2*. Each curve represents the average of three biological triplicates each with 4 technical replicates, with the shade showing one standard deviation. f-g) Maximum projection images of example z-stacks showing f) FRQ or g) WC-1 subcellular localization (left) and quantification of nuclear signal (right) in constant light, with or without Impβ2. Scale bar = 10 μm. N = 3-6 hyphal tips. Each dot represents the mean fluorescent signal of one nucleus. h) Nuclear FRQ signal recovery speed at CT4 with or without Impβ2, calculated as the slope of linear regression of each target nucleus’s recovery curve.

A full knockout of *imp*β*2* causes minor growth defects (Fig. S4f), shortens circadian period by around 2.5 hours, and slightly reduces WCC activity leading to loss of persistence of the rhythm (Fig. 5d-e). Consistent with this, knockdown or knockout of TNPO1 in mammalian cells also shortens circadian period (Korge et al. 2018), suggesting a similar regulatory mechanism between the two circadian systems. However, nuclear localization of neither FRQ nor WC-2 is impaired in the Δ*imp*β knockout strains (Fig. 5f-g). FRAP experiments at CT4 showed no difference in FRQ nuclear import rate either (Fig. 5h). There are also no obvious changes in core clock protein expression levels in the Δ*imp*β knockout strains (Fig. S4g).

*imp*β*3* is also essential, but viable conditional knockdowns were achieved by insertion of the inducible *qa-2* promoter directly upstream of the ORF (Fig. 6a). A VHF tag was fused to its C terminus to facilitate protein detection. In the absence of QA, the expression level of Impβ3 is significantly lower compared to QA-induced levels (Fig. 6b, Fig. S5a), resulting in a substantial circadian period length shortening to around 15 hours (Fig. 6c). A relatively small amount of Impβ3 is sufficient to sustain a normal clock, as even a low concentration (10^-4^M) of QA restores the period to wild-type length, and higher QA concentrations have no additional effect (Fig. 6c, Fig. S5b-c). Expression of core clock proteins remains normal in Impβ3-deficient samples (Fig. 6b, Fig. S5a). To examine subcellular localization of core clock components, endogenous fluorescent-tagged *frq* or *wc-2* alleles were introduced into the inducible strain, which exhibited dramatically shortened periods as expected (Fig. S5d). Despite the period effects, loss of Impβ3 resulted in no reduction in nuclear localization of either FRQ or WC-2 (Fig. 6d-e).

**Figure 6.**
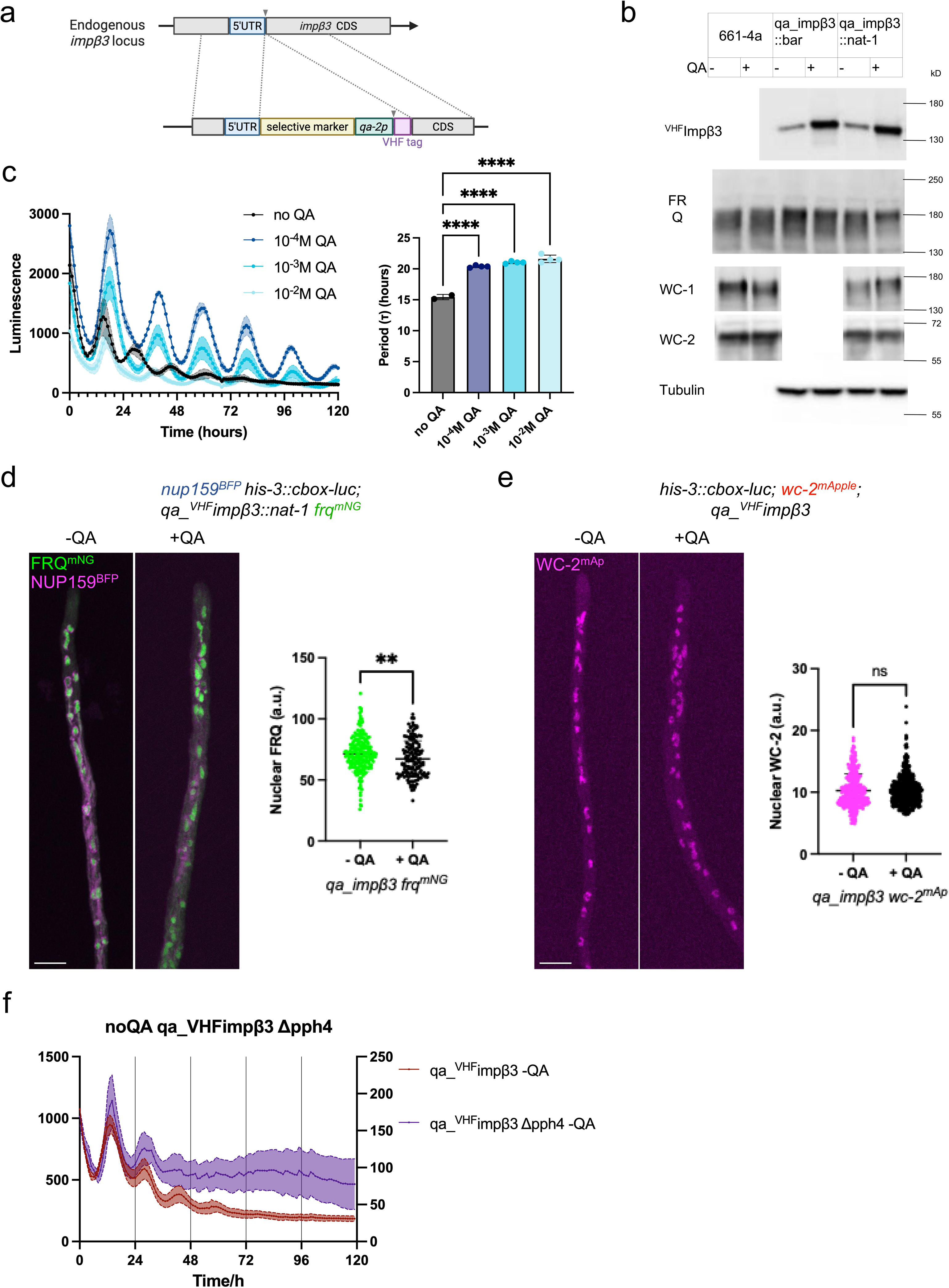
Shortened period in Impβ3-deficient strains is independent of core clock protein levels or localization. a) Schematic of the inducible *imp*β*3* design. Expression of *imp*β*3* is driven by the *qa-2* inducible promoter. Without the addition of the inducer quinic acid (QA), expression of *imp*β*3* is reduced. b) Core clock protein levels in the control strain or inducible *imp*β*3* strains with or without 10^-2^ M QA. The result shown is representative of biological triplicates; additional replicates are shown in Fig. S5d. c) Luciferase assays of the inducible *imp*β*3* strain with varying QA concentrations (left) and their period lengths (right). n=4 replicates. The shade shows one standard deviation. Periods were analyzed by one-way ANOVA with multiple comparisons. d-e) Maximum projection images of example z-stacks showing d) FRQ or e) WC-2 subcellular localization ((left) and quantification of nuclear signal (right) in constant light, with or without 10^-2^ M QA. Scale bar = 10 μm. N = 4-7 hyphal tips. Each dot represents the mean fluorescent signal of one nucleus. Two sibling strains were quantified for WC-2. f) Luciferase assays of strains with both *pph-4* deletion and *imp*β*3* deficiency compared to strains with *imp*β*3 deficiency* alone. n=4 replicates. The shading shows one standard deviation.

Period shortening, particularly the dramatic period shortening seen with reduced Impβ3, is less often seen with clock mutants. This prompted us to further investigate the possible mechanism by which Impβ3 regulates the circadian clock by looking for possible interactions with protein phosphatase 4 (PPH-4; also known as PP4, NCU08301) whose loss also results in period shortening (Cha et al. 2008). PPH-4 has previously been shown to regulate the nuclear enrichment of the WCC, and loss of *pph-4* also results in a shortened circadian period, similarly to Impβ3 knockdown (Fig. S5e) (Cha et al. 2008; Dasgupta 2015). Interestingly, the period phenotypes caused Δ*pph4* and *imp*β*3* reduction are not additive; that is, knockdown of *imp*β*3* is epistatic to *pph4*. When Impβ3 is present (with QA), Δ*pph-4* caused the expected period shortening (Fig. S5e). However, using the same pair of strains, combining Δ*pph-4* with Impβ3 reduction (no QA) does not further shorten the circadian period although rhythm persistence is reduced (Fig. 6f, Fig. S5e). This non-additive effect suggests that Impβ3 and PPH-4 act within the same or overlapping pathways to regulate circadian rhythmicity. Impβ3 may facilitate the proper subcellular localization of PP4 or its associated co-enzymes, thereby enabling PP4 to precisely regulate phosphorylations of core clock proteins. These findings point to functional crosstalk between nucleocytoplasmic transportation and phosphatase-mediated regulation in maintaining the accuracy of circadian clock.

## DISCUSSION

Our findings establish nuclear import as a key regulatory step in the *Neurospora* circadian clock. Using the live-cell imaging platform (Wang et al. 2026), we are able to measure in vivo subcellular dynamics in *N. crassa* under native, biologically relevant conditions. Through FRAP, we observed that FRQ is imported into nuclei slowly during the subjective day, requiring >20 min to recover fluorescence (Fig. 1). This is in contrast with an earlier report of rapid nucleocytoplasmic shuttling of FRQ based on an anchor-away strategy that fused FRQ to FRB and tethered it to FKBP-tagged histone H1 (Diernfellner et al., 2009). While powerful, that system relies on rapamycin-induced heterodimerization of FRB-FKBP, which introduces an artificial import route that could diminish intrinsic clock function, raising the possibility that the observed fast shuttling is a result of methodological differences rather than the intrinsic behavior of FRQ in a free-running circadian cycle.

FRAP analyses revealed that FRQ nuclear import is not constant but varies across different stages of FRQ nuclear accumulation of the circadian cycle, being most active early in the subjective day and strongly weakened near subjective dusk (Fig. 1). This temporal regulation ensures that nuclear FRQ accumulation occurs with proper timing and rate, contributing to period stability. A previous report using FRAP to estimate nuclear import of mammalian Negative Arm proteins tested only extreme time points - when nuclear levels were at their peak or trough (Smyllie et al. 2016). As these conditions represent equilibrium states with little net nucleocytoplasmic exchange, it may be reasonable that no difference in import rate was detected (Smyllie et al. 2016). In contrast, by assaying import continuously across the subjective day, we observed a gradual decrease in nuclear import as FRQ approached its peak nuclear accumulation. The timescale of FRQ import we observed by FRAP (tens of minutes for recovery) is comparable to that of PER1 and PER2, functional homologs of FRQ in mammalian systems, whether assayed through endogenous tagging or transient expression (Öllinger et al. 2014; Smyllie et al. 2016; Korge et al. 2018). This similarity across fungal and mammalian systems suggests that slow nuclear import is a conserved feature of Negative Arm proteins and may represent an important delay step required to sustain a ∼24-hour feedback loop.

Impα has been shown to regulate the subcellular localization of the Negative Arm in both mammalian systems and *Drosophila* (Umemura et al. 2014; Jang et al. 2015). The dependence of FRQ nuclear import on Impα, supported by both biochemical Co-IP and fluorescent imaging after RNAi knockdown experiments, places Impα at the core of this regulatory process (Fig. 2-4). Worth noting is that the FRQ bound by Impα is hypophosphorylated, consistent with previous research showing that newly synthesized FRQ tends to accumulate in nuclei (Luo et al. 1998; Hong et al. 2008; Diernfellner et al. 2009) and supporting a model in which phosphorylation state modulates FRQ nuclear import efficiency. Future investigation combining FRAP and phospho-specific analysis may further resolve the detailed regulatory mechanism. Impα facilitates nuclear import by directly binding to the NLS on a target protein (Conti et al. 1998). Surprisingly, a domain previously shown to be essential for circadian rhythmicity and suggested to function as a NLS on FRQ (Luo et al. 1998), is not required for nuclear localization of FRQ (Fig. 2) (Schunke et al. 2026). These strains exhibit persistently high WCC activity, suggesting that the loss of rhythmicity arises from a failure of FRQ to inhibit WCC despite its nuclear presence. Deletion of this motif has also been proposed to disrupt FRQ dimerization (Lauinger et al. 2014), which is essential for the circadian clock and could be the cause of arrhythmicity (Cheng et al. 2001). Very recent work overexpressing fluorescent tagged FRQ fragments in mammalian cell culture support our conclusion and suggest the possible presence of multiple redundant NLSs on FRQ (Schunke et al. 2026). Similar redundancy has also been reported for the Positive Arm WCC, whose nuclear localization doesn’t rely on a single NLS (Schwerdtfeger and Linden 2000; Cheng et al. 2003; Wang et al. 2016). Together, these observations suggest that nuclear import in *Neurospora* is governed by a more complex and possibly redundant regulatory mechanism compared to that seen in other eukaryotes (Sakakida et al. 2005; Kwon et al. 2006; Saez et al. 2011), perhaps consistent with the unique challenges posed by its syncytial organization.

In addition to FRQ, our results demonstrate that WC-1 nuclear import is also strongly dependent on Impα (Fig. 3-4). In Impα-deficient strains, WC-1 protein became undetectable despite normal transcript levels, consistent with post-transcriptional destabilization. It may be that WC-1 is unstable in the cytoplasm and is rapidly degraded when nuclear import is blocked, thereby explaining the dramatic loss of WC-1 protein in Impα-deficient strains. By contrast, WC-2 nuclear import, when on its own, does not require Impα and its local level even increases with reduced Impα. This increase is possibly an indirect effect through loss of WC-1, as WC-1 has been proposed to be a repressor of *wc-2*: in Δ*wc-1*, both *wc-2* RNA and WC-2 protein levels are high compared to wildtype (Cheng et al. 2003; Neiss et al. 2008). Although the overall amount of nuclear WCC does not show strong oscillations (Denault et al. 2001), WCC has been reported to undergo rapid nucleocytoplasmic shuttling (Schafmeier et al. 2008). NCU01249 encodes the sole Impα subunit in *Neurospora*, which must control nuclear import of numerous cargo proteins and has been linked to additional cellular functions beyond nuclear import (Klocko et al. 2015). Therefore, the effects of Impα deficiency on both growth and circadian rhythmicity is likely mediated through multiple pathways beyond altered localization of core clock proteins.

Our analysis of Impβ proteins further underscores the complexity of importin-mediated regulation (Fig. 5-6). Impβ1 is essential and difficult to genetically manipulate, but chemical inhibition suggests a role in FRQ nuclear localization, possibly in cooperation with Impα. Impβ2 deletion shortened the circadian period by ∼2.5 h, paralleling results from knockdowns of its mammalian homolog TNPO1 (Korge et al. 2018). However, nuclear localization of FRQ and WC-2 was unaffected, suggesting that Impβ2 influences the clock through other cargoes or regulatory pathways. Impβ3 is also important for precise circadian control: reduced expression caused dramatic period shortening without altering nuclear localization of FRQ or WC-2, again suggesting other cargoes or regulatory pathways. Genetic interaction experiments with *pph-4* indicate that Impβ3 and PPH4 act within the same regulatory pathway, with Impβ3 potentially functioning upstream to facilitate PPH4 activity or localization. This unexpected connection highlights the interplay between nuclear transport and phosphatase-mediated regulation of WCC.

Together, these results demonstrate that nuclear import is a dynamic and selective process that plays a critical role in shaping circadian timing in *Neurospora*. Our findings parallel observations in other eukaryotic systems (Sakakida et al. 2005; Saez et al. 2011; Öllinger et al. 2014; Umemura et al. 2014; Jang et al. 2015; Lee et al. 2015; Korge et al. 2018; Korge et al. 2018), underscoring the evolutionary conservation of importin-mediated circadian regulation, while also revealing unique fungal-specific mechanisms.

## MATERIALS AND METHODS

### Strain construction

Plasmids were constructed using pRS426 as backbone. Gene fragments were amplified from genomic DNA of *N. crassa* 74-OR23-1VA (FGSC, 2489) using Phusion Flash High-Fidelity PCR Master Mix (Thermo Scientific # F548S). Other fragments were either ordered as a gBlock or amplified from plasmids previously generated in the lab. Fragments were then assembled together using NEBuilder® HiFi DNA Assembly Master Mix (New England Biolabs Cat. #E2621) according to the manufacturer’s instructions. All plasmids and their sequence data have been deposited at Addgene.

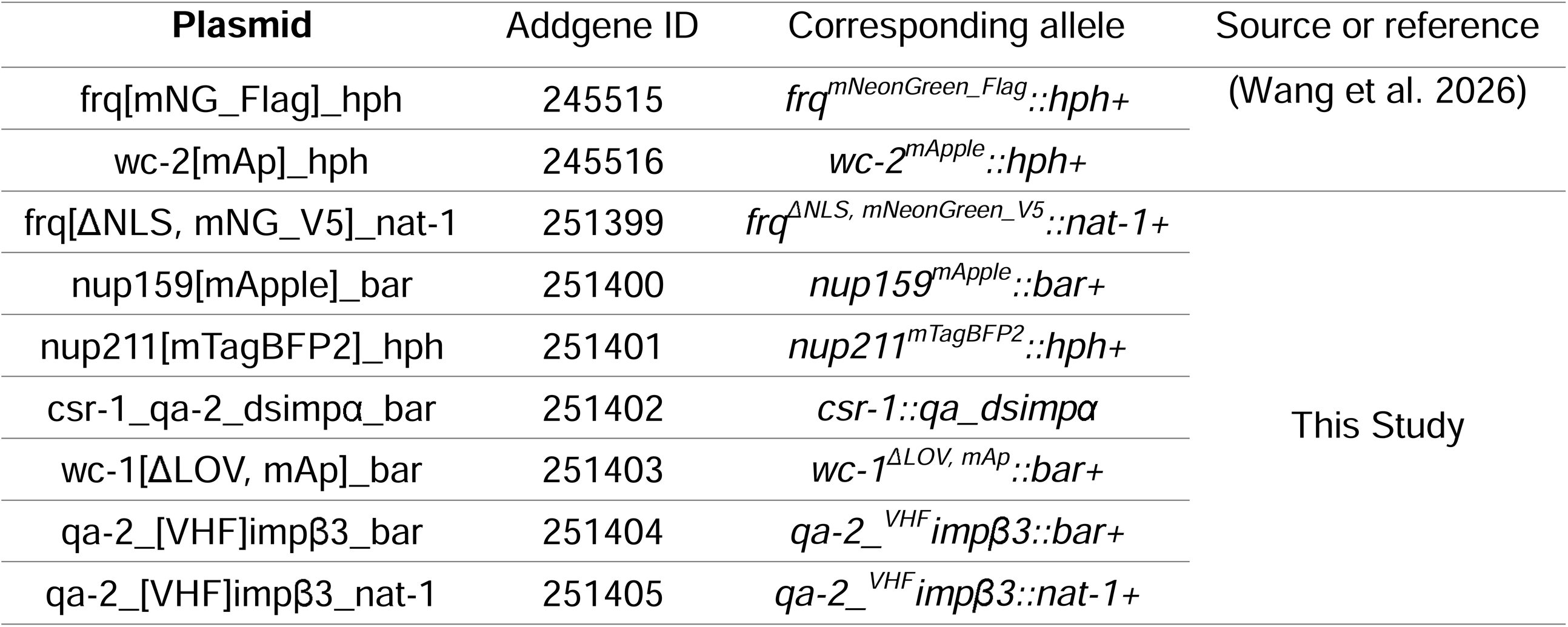

*frq*^Δ*NLS,*^ *^mNeonGreen_V5^::nat-1+* contains the full *frq* gene with NLS1 (Luo et al. 1998) deleted, fused at its C-terminus to codon-optimized mNeonGreen and a V5 tag, followed by the nourseothricin resistance marker (He et al. 2020), designed to target the endogenous *frq* locus. The transformation cassette was amplified using Phusion Flash High-Fidelity PCR Master Mix (Thermo Scientific # F548S) and then transformed into strain 576-4 (Δ*mus-52::hph+;* Δ*frq::hph+*).

*nup159* (NCU08604) and *nup211* (NCU04059) were endogenously tagged at their C-termini using similar strategy as described previously (Bartholomai 2022) using different fluorescent tags and drug selection markers. *imp*α (NCU01249) was endogenously V5-tagged at its C-terminus using a hygromycin resistant cassette for selection, resulting in the *imp*α*^V5^::hph+* allele.

The expression cassette producing double-stranded RNA targeting exon 3 of *importin* α (NCU01249) under control of the QA-inducible promoter was constructed as shown in Fig. 3a and inserted at the *csr-1* locus.

The *qa-2* promoter, along with a V5_6xHis_3xFlag (VHF) tag, was inserted at the N-terminus of *importin* β*3* (NCU03257) coding sequence using standard lab protocols to regulate its expression (Aronson et al. 1994). Two series of strains with different drug selection markers were constructed with other alleles during sexual crosses.

Transformations of *Neurospora crassa* were performed following lab protocols as described previously (Bartholomai 2022), after which sexual crosses were carried out using standard Neurospora methods (http://www.fgsc.net/Neurospora/NeurosporaProtocolGuide.htm) to generate homokaryotic progeny with the desired combination of alleles.

### FRAP

5 independent runs were performed for FRAP experiments using strain 1946-1 (*csr-1::son-1^mTagBFP2^; ras-1^bd^; wc-1*^Δ*LOV*^*::bar+, for, frq^mNeonGreen_Flag^::hph+*) in the customized microfluidic devices as described previously (Wang et al. 2026) with Bird medium (Metzenberg 2004) containing 0.2% glucose, 0.2% poly-acrylic acid, and 1mM sodium formate. The cultures were entrained by temperature steps before resetting to CT0. 2.5 μg/ml benomyl was added to the medium at the same time as resetting. For each independent run, one FRAP experiment was performed on a new region within the same culture every half an hour from CT2 to CT10. Single focal plane images were acquired on LSM 880 with a Plan-Apochromat 63x/1.4 oil objective and with the pinhole size adjusted to an optical section of 2.8 μm. For each recording, 40 to 50 frames were collected at 30-second intervals. After the first two frames, 4-7 nuclei were photobleached with a 488 nm laser to around 30% of the initial fluorescence intensity. At CT2, low nuclear FRQ levels (Wang et al. 2026) limited effective photobleaching, so fewer nuclei met the criteria to be included in later analysis. SON-1^mTagBFP2^ (DAPI channel) was recorded at the same time as nuclear marker.

Measurements of F_ROI (manually bleached nuclei), P_ROI (untreated nuclei) and B_ROI (outside of the cell) were taken in ImageJ. Rolling averages of 3 time points (n-1, n, n+1) were calculated and used for further analysis. Fluorescence recordings were corrected for background. The fitted values based on an averaged slope (photobleaching rate) calculated from the linear regressions of *P_ROI - B_ROI* within the same run were used to correct for photobleaching during recording:

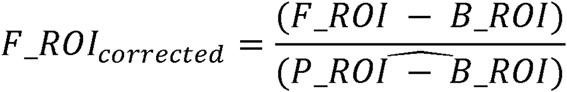

*F_ROI_corrected_* was then normalized to pre-beach intensities (set as 100%) and grouped into 1-hour bins, i.e. all *F_ROI_corrected_*from HT3 and HT3.5 were considered as group HT3, for Fig. 1c.

A linear regression was calculated for each nucleus (*F_ROI_corrected_*). Then, the slopes of these fitted curves were grouped into 1-hour bins for Fig. 1d. A small number of nuclei were excluded

| Time | CT2 | CT3 | CT4 | CT5 | CT6 | CT7 | CT8 | CT9 | CT10 |
| --- | --- | --- | --- | --- | --- | --- | --- | --- | --- |
| N of included nuclei ($r^2 \geq 0.5$) | 4 | 22 | 26 | 32 | 35 | 37 | 32 | 24 | 18 |
| N of excluded nuclei ($r^2 < 0.5$) | 0 | 1 | 1 | 1 | 0 | 1 | 1 | 2 | 0 |

because of poor goodness of fit ( $r^2$ ).

For Fig. S1g, a similar analysis was performed using linear regression of a single example *P_ROI - B_ROI* within the same FRAP experiment (same recording/region) to correct for photobleaching, rather than using the average in one run. Recordings without a suitable P_ROI were excluded from the analysis.

| Time | CT2 | CT3 | CT4 | CT5 | CT6 | CT7 | CT8 | CT9 | CT10 |
| --- | --- | --- | --- | --- | --- | --- | --- | --- | --- |
| N of included nuclei ($r^2 \geq 0.5$) | 4 | 21 | 20 | 25 | 26 | 36 | 28 | 24 | 16 |
| N of excluded nuclei ($r^2 < 0.5$) | 0 | 0 | 0 | 0 | 1 | 2 | 1 | 2 | 2 |

were excluded from the analysis.

### Luciferase assay

All luciferase assays were performed and analyzed in the same way as described previously (Kelliher et al. 2023). Unless otherwise specified, all entrainments and recordings were performed at 25°C. For the benomyl test in Fig. S1d, 50 μl per well of water or 2.5 μg/ml benomyl was injected at the indicated times using 1ml syringes. All other drugs, as well as the inducer quinic acid, were added directly to the medium before inoculation at the indicated concentrations.

### Immunoprecipitation and Western blot

Co-IP experiments were performed following previously described protocols with slight modifications (Chen et al. 2010). Protein samples were collected, and Input samples were saved, using the same method described in Wang et al., (2025). All protein samples were from whole cell lysates. For V5 IP, V5 beads were prepared by preincubating 2μl V5 antibody (Invitrogen #46-0705) with 30μl Dynabeads^TM^ Protein G (Invitrogen #1003D) per sample at 4°C for 1 hour. After determining protein concentrations in the protein samples by Bradford assay, 2mg protein per sample in a 1ml volume was incubated with the V5 beads at 4°C for 2 hours or overnight. The bound complex was then eluted with 100μl 2x Sample Buffer (diluted from NuPAGE™ LDS Sample Buffer (4X), Invitrogen #NP0007) following two washes with Wash Buffer (50mM Hepes-KOH, pH 7.4, 150mM NaCl, 0.4% NP-40) and two washes with no-NP Wash Buffer (50mM Hepes-KOH, pH 7.4, 150mM NaCl). For detection, 30μg Input and 10μl IP (from approximately 200μg total protein) samples were loaded onto NuPAGE™ 3-8% Tris-Acetate Mini Protein Gels (ThermoFisher Scientific, Catalog #EA03785BOX). Western blotting was then performed as described in Wang et al., (2026).

For the *qa_dsimp*α samples, cultures were initiated in 10cm dishes with liquid culture medium (LCM) (1× Vogel’s salts, 2% glucose, 0.5% arginine, 50 ng/mL biotin) without QA. Five ∼1cm diameter circles were cut and transferred to 275ml flasks containing 50ml LCM, with or without 10^-2^M QA (as indicated in figures), and cultured for 21 hours before harvest. For all other samples, strains were maintained in the same LCM (with or without 10^-2^M QA) throughout the culture. All cultures were performed in light at 25°C unless stated otherwise.

### Fluorescent imaging

Samples were grown at 25 °C in the light on agarose pads (1× Vogel’s salts, 2% sucrose, 0.5% arginine, 50 ng/mL biotin, 1% agarose) overnight, unless otherwise specified, and imaged using the ‘inverted agar block’ method (Hickey et al. 2004) on one of two Nikon Eclipse Ti-E stand-based spinning disk confocal microscope systems. Each microscope is equipped with a Nikon LU-N4 laser launch that includes 405 nm, 488 nm, 561 nm, and 640 nm lasers , a Yokogawa CSU-W1 spinning disk system, two Photometrics Prime BSI sCMOS cameras, an ASI MS-2000 motorized stage with a piezo Z-drive, and a Tokai Hit stage-top incubation system for temperature control. All images were acquired with a Nikon CFI Plan Apochromat λD 60× oil objective with a numerical aperture of 1.42.10 μm z-stacks (35 slices at 300 nm step size) were collected in corresponding channels. Tip regions were cropped in ImageJ and then 3D reconstituted in Arivis Vision 4D for segmentation and quantification. Mean fluorescent signals from both the total hyphae and individual nuclei were measured and background signals, defined as the mean intensity from a manually generated object outside the hyphae in each z-stack, were subtracted. The coefficient of variation (CV) for each nucleus was calculated as the standard deviation of its fluorescence signal (SD) divided by its background subtracted mean fluorescent signal (MEAN_background substracted_):

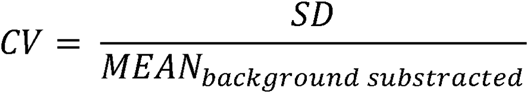

For DD samples in Fig. 2c-d and Fig. 5a-c, samples were germinated on agarose pads (without drug) in light and then transferred to darkness for the indicated number of hours. For Fig. 5a-c, at DD12, a small piece of agarose at the growth front was cut, and its top surface was immersed in water or a 40μM drug solution. After 2 hours of drug and light exposure, samples were imaged using the same methods. Quinic acid was added to the agarose pad prior to inoculation in corresponding experiments.

### Race tubes / growth tests

Race tube assays were performed as described previously (Aronson et al. 1994). For growth tests with Impβ1 inhibitors, race tube medium was supplemented with the indicated concentrations of drug. Conidia of strain 661-4a (*his-3::frq_cbox_-luc*) were inoculated at the edge of 35-mm dishes and incubated overnight at 25 °C in constant light before being scanned.

### RT-qPCR

Samples were cultured and harvested as described for Western blotting. RNA was extracted from ∼200μg ground powder per sample using Direct-zol RNA Miniprep Kits (ZYMO Research, R2050). For cDNA synthesis, 3μg extracted RNA reverse-transcribed using SuperScript™ IV First-Strand Synthesis System (Invitrogen, 18091050) with the provided Oligo d(T)_20_ primer. The resulting cDNA was diluted to ∼7.5 ng/μl, and 2 μl was used for RT-qPCR with iTaq Universal SYBR Green Supermix (Bio-Rad, #1725121) measured with an Applied Biosystems™ StepOne™ Real-Time PCR System. The primers used for RT-qPCR are listed below:

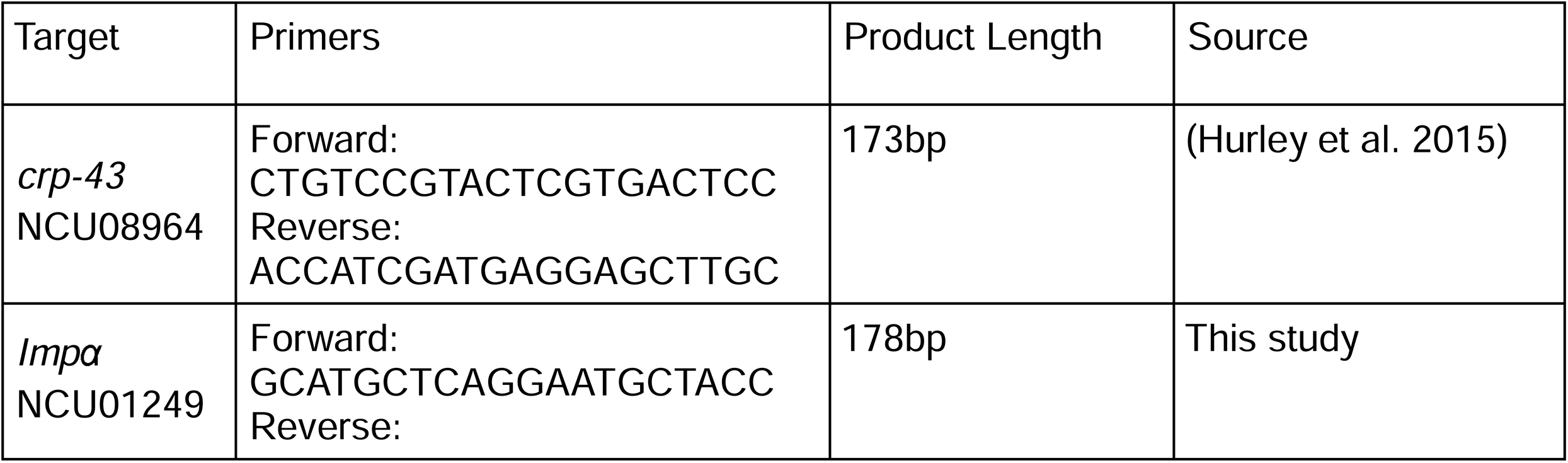
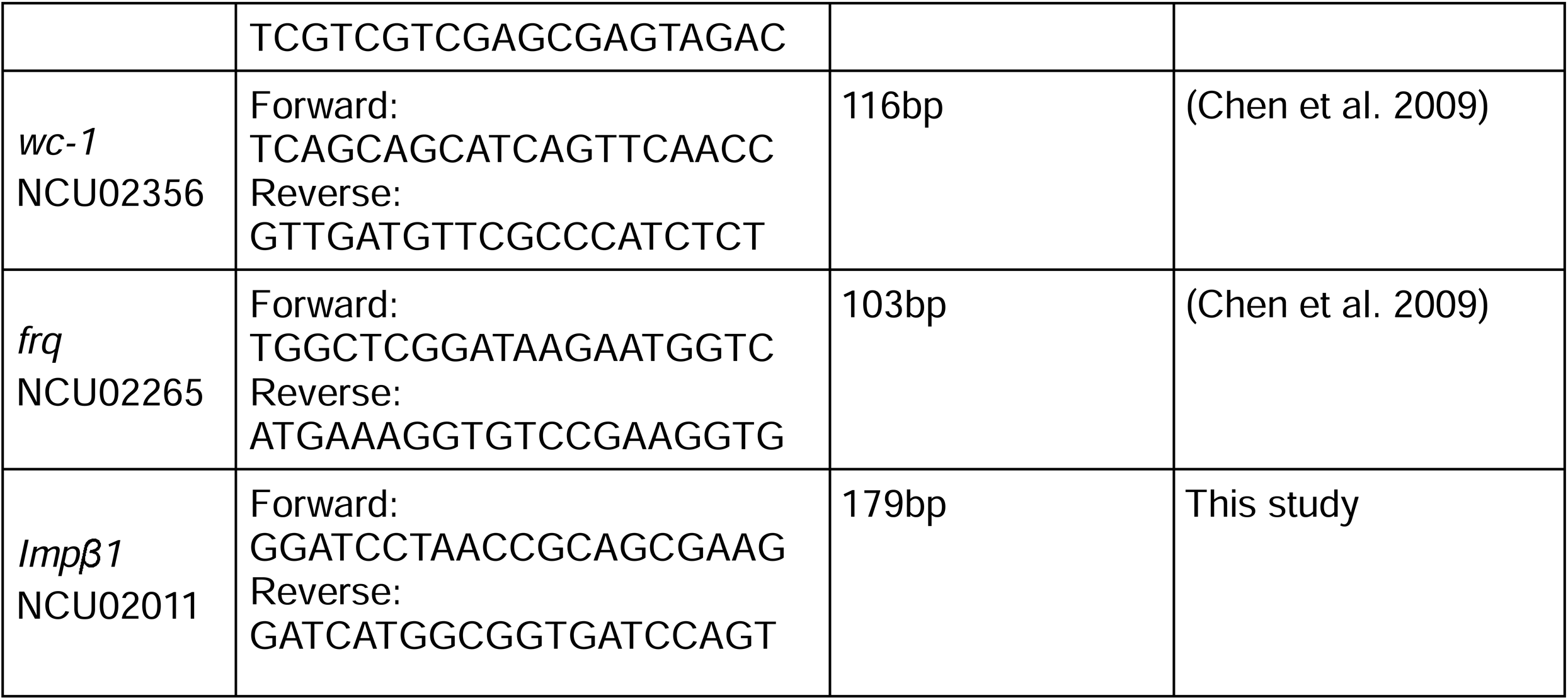

For light pulse experiments, samples were initiated in light, the cut circles were transferred to flasks in the dark and cultured for 21□h. Cultures were then either harvested in dark (DD21) or exposed to light for 15 minutes before harvest (15min LP).

### Statistical analysis and data visualization

Statistical analysis was performed using Prism 10 (GraphPad) with either one-way ANOVA followed by multiple comparisons or Welch’s t test. The number of replicates (n) used for each experiment are indicated in the relevant figure legends. All schematic diagrams were created in BioRender. Wang, Z. (2026) https://BioRender.com/8cxiutk.

## DATA AVAILABILITY STATEMENT

The authors affirm that all data necessary for confirming the conclusions of the article are present within the article, figures, and tables. Strains are available upon request and all plasmids have been deposited in and are available from Addgene.

## ACKNOWLEDGMENTS

This study was supported by a National Institutes of Health (NIH) grant awarded to Jay C. Dunlap (R35GM118021). We acknowledge use of stocks from the Fungal Genetics Stock Center (FGSC.net), NIAID-supported informatic resources at FungiDB (Contract HHSN75N93019C00077), and Molecular Biology facilities supported by the Dartmouth CQB COBRE (NIH NIGMS grant P20GM130454), and the Dartmouth BioMT (NIH NIGMS grant P20-GM113132). Imaging was performed in the Life Sciences Light Microscopy Facility (RRID:SCR_027143) at Dartmouth College.

