## Supplementary Materials for "Rhythmic Nuclear Import Mediated by Importins Regulates the *Neurospora* Circadian Clock"

**Supplementary Table 1 | Potential nuclear transporters in *Neurospora crassa*.**

| *Neurospora crassa* | *Saccharomyces cerevisiae* | *Aspergillus nidulans* | Human |
|---|---|---|---|
| **NCU01247 Importin subunit alpha** | Kap60p/SRP1 [YNL189W] | AN2142 kapA | |
| **NCU02011 importin subunit beta-1** | Kap95p [YLR347C] | AN0906 kapB | KPNB1 (karyopherin subunit beta 1) |
| **NCU03690 importin beta-2 subunit** | Kap104p [YBR017C] | AN0926 kapC | TNPO1 (transportin 1) and TNPO2 (transportin 2) |
| NCU06355 Karyopherin | Kap111p/Mtr10p [YNL153C] | AN6734 kapF | TNPO3 transportin 3 |
| NCU06578 KapG | Kap114p [YGL241W] | AN2164 kapG | IPO9 (importin 9) |
| NCU16337 nucleoporin-9 | Kap120p/Lph2p [YPL125W] | AN4053 kapH | IPO11 (importin 11) |
| **NCU02357 importin subunit beta-3** | Kap121p/Pse1p [YMR308C] | AN5717 kapI | several human genes including IPO5 (importin 5) |
| NCU16656 hypothetical protein | Kap122p/Pdr6p [YMR192W] | AN7731 kapN | IPO13 (importin 13) |
| NCU05650 karyopherin Kap123 | Kap123p/Yrb4 [YER110C] | AN2120 kapJ | IPO4 (importin 4) |
| NCU01820 exportin-1 | Kap124p/Crm1p/Xpo1p [YGR218W] | AN1401 kapK | Exportin 1 (CRM1, XPO1) |
| NCU04104 chromosome segregation protein Cse1 | Cse1p/Kap109p [YGL137W] | AN6591 kapE | CSE1L (chromosome segregation 1 like) |
| NCU00134 exportin-T | Los1p [YPL084W] | AN8787 kapM | XPOT (exportin for tRNA) |
| NCU02387 nuclear import and export protein Msn5 | Kap142p/Msn5p [YDR335W] | AN3012 kapL | XPO5 (exportin 5) |

*NCU01939 is identified as a weak homolog of both Kap108p/SXM1 [YPL147W] and Kap119p/Nmd5p [YGL226W] by OthoDB, but not by NCBI Orthologs.

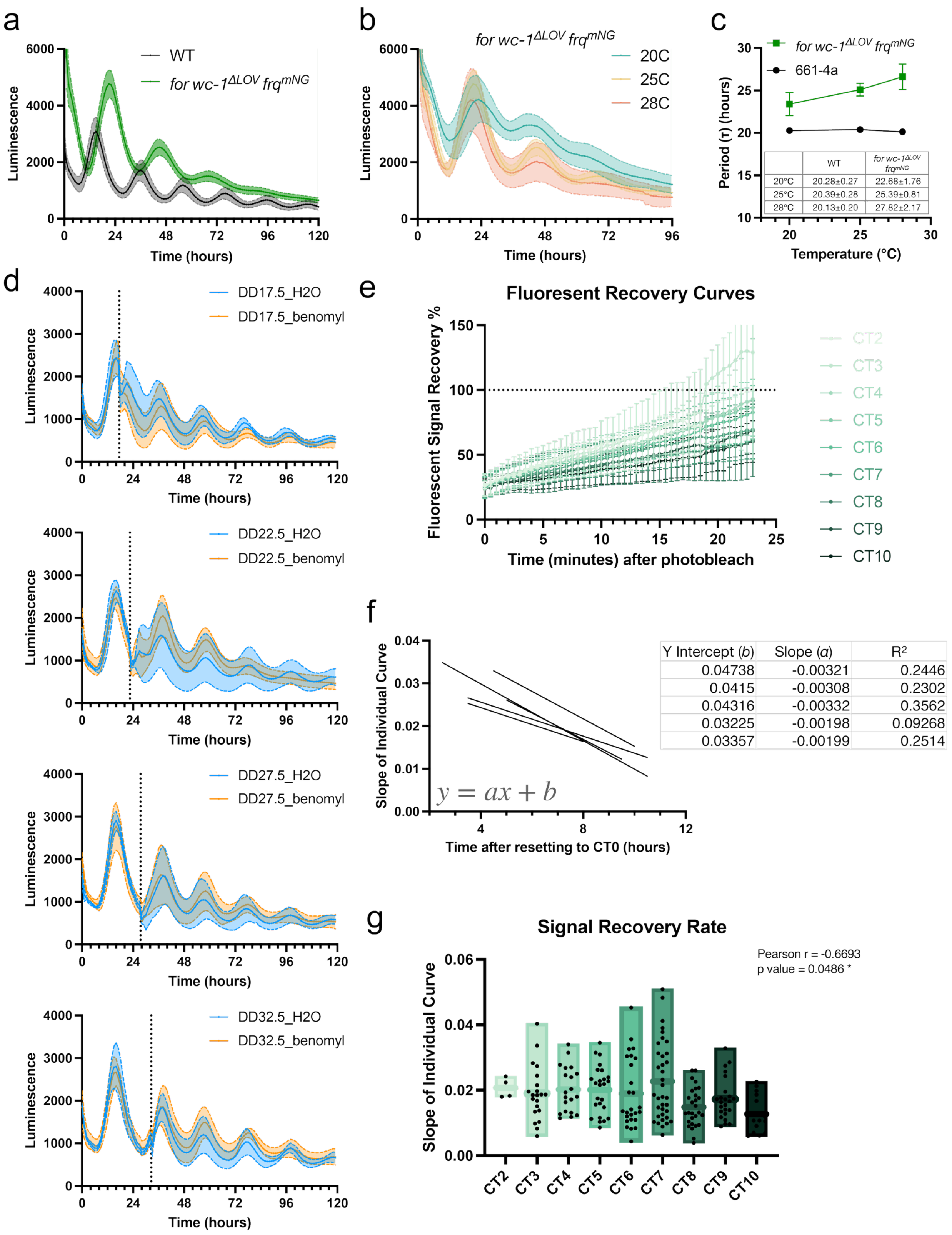
a
Luminescence
Time (hours)
WT
for wc-1ΔLOV frqmNG
b
for wc-1ΔLOV frqmNG
20C
25C
28C
Luminescence
Time (hours)
c
for wc-1ΔLOV frqmNG
661-4a
Period (τ) (hours)
Temperature (°C)
WT
for wc-1ΔLOV frqmNG
20°C 20.28±0.27 22.68±1.76
25°C 20.39±0.28 25.39±0.81
28°C 20.13±0.20 27.82±2.17
d
DD17.5_H2O
DD17.5_benomyl
DD22.5_H2O
DD22.5_benomyl
DD27.5_H2O
DD27.5_benomyl
DD32.5_H2O
DD32.5_benomyl
Luminescence
Time (hours)
e
Fluoresent Recovery Curves
Fluorescent Signal Recovery %
Time (minutes) after photobleach
CT2
CT3
CT4
CT5
CT6
CT7
CT8
CT9
CT10
f
Slope of Individual Curve
Time after resetting to CT0 (hours)
y = ax + b
Y Intercept (b) Slope (a) R2
0.04738 -0.00321 0.2446
0.0415 -0.00308 0.2302
0.04316 -0.00332 0.3562
0.03225 -0.00198 0.09268
0.03357 -0.00199 0.2514
g
Signal Recovery Rate
Pearson r = -0.6693
p value = 0.0486 *
Slope of Individual Curve
CT2
CT3
CT4
CT5
CT6
CT7
CT8
CT9
CT10

**Supplementary Figure 1 | Optimized FRAP experiments elucidate circadian regulated nuclear import of FRQ.**

a-b) Luciferase assays of the dual-color imaging strain *his-3::frq*$_{cbox}$*-luc, ras-1*$^{bd}$*, wc-1*$^{\Delta LOV}$ *for frq*$^{mNeonGreen}$ (1952-2) (a) at 25℃ compared to wildtype or (b) across temperatures. Each curve represents the average of three biological triplicates each with 4 technical replicates, with the shade showing standard deviations. c) Periods calculated from luciferase assays of the imaging strain (1952-2) at different temperatures after temperature entrainments compared to wildtype. n=12 replicates for each condition. Error bars show standard deviations. d) Luciferase assays showing the effect of benomyl addition, compared to water, at different circadian phases. Each curve represents mean ± SD from n=4 replicates. e) Average fluorescent recovery curves, with variations (standard deviation) for each hour of the subjective day. f) Linear regression of nuclear signal recovery speed across time for each individual time course (N = 5 time courses). The parameters for each linear regression are included in the table. g) Nuclear signal recovery speed, calculated as the slope of linear regression of the recovery curve for each target nucleus, when using a control nucleus within the same recording for photobleaching correction. Data were analyzed by Pearson correlation, with the correlation coefficient (r) and p-value indicated in the plot.

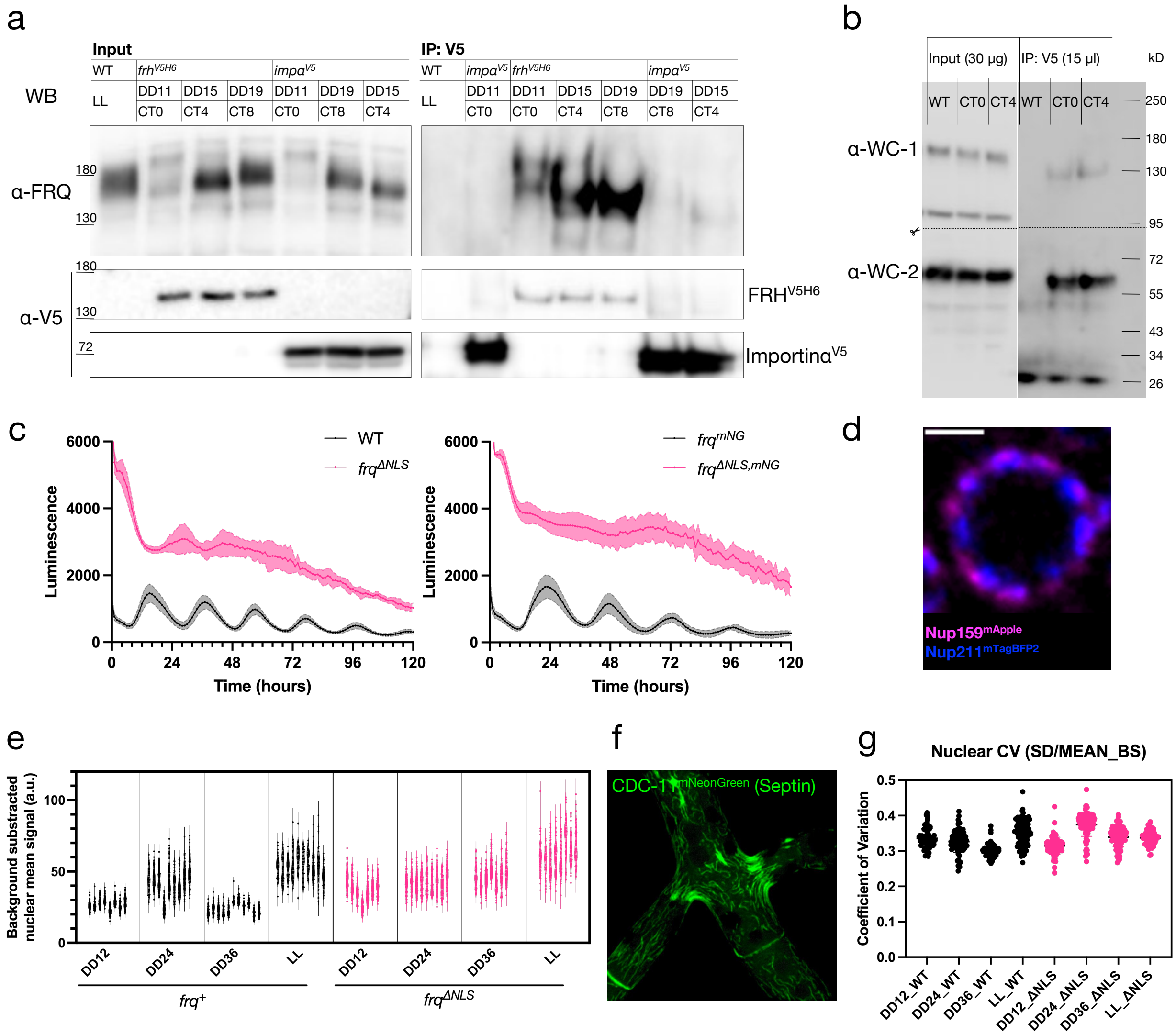


**Supplementary Figure 2 | Rhythmic nuclear oscillation of FRQ is abolished in *frq*$^{\Delta NLS}$.**

a-b) Biological replicates of co-immunoprecipitation using anti-V5 antibody showing the interaction between Importin α and a) FRQ or b) WCC. Following transfer, PVDF membranes were cut (dotted line) according to expected protein sizes and probed with the indicated antibodies. c) Luciferase assays of *frq*$^{\Delta NLS}$ strains compared to their corresponding control strains with wildtype *frq*. Each curve represents mean ± SD from n=4 replicates. d) Super resolution image of the inner nuclear envelope marker (NUP211 in blue) and the outer nuclear envelope marker (NUP159 in magenta) acquired using SoRa system. Scale bar = 1μm. e) Nested violin plot showing ungrouped nuclear FRQ or FRQ$^{\Delta NLS}$ signal from individual hyphal tips. f) Example image of CDC-11 (septin) endogenously tagged by mNeonGreen. g) Subnuclear heterogeneity of FRQ or FRQ$^{\Delta NLS}$, represented by coefficient of variation, during free-run circadian cycles or in constant light.

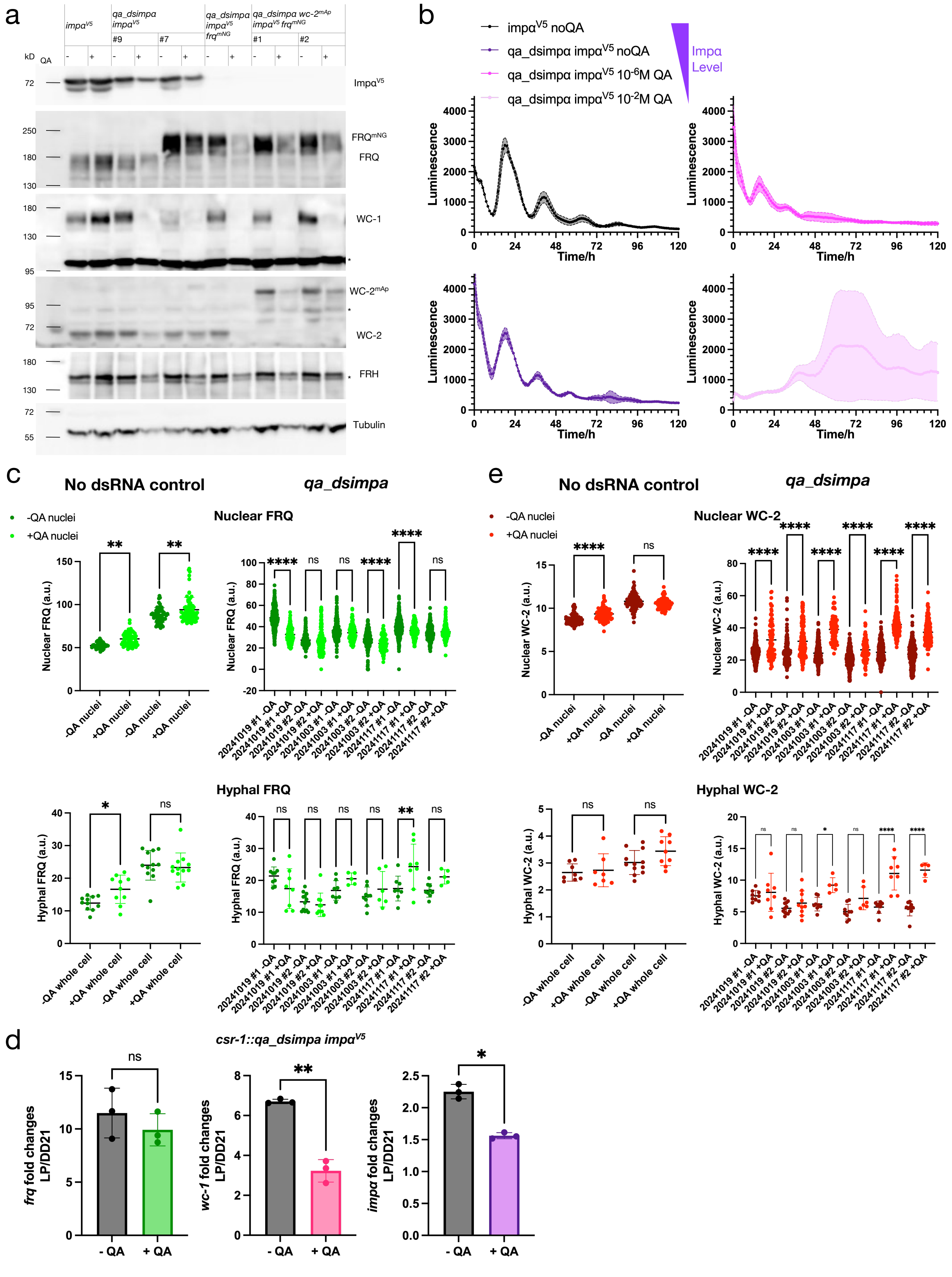
a
impα^V5
qa_dsimpa impα^V5
qa_dsimpa impα^V5 frq^mNG
qa_dsimpa wc-2^mAp impα^V5 frq^mNG
#9
#7
#1
#2
kD
QA
Impα^V5
FRQ^mNG
FRQ
WC-1
WC-2^mAp
WC-2
FRH
Tubulin
b
impα^V5 noQA
qa_dsimpα impα^V5 noQA
qa_dsimpα impα^V5 10^-6M QA
qa_dsimpα impα^V5 10^-2M QA
Impα Level
Luminescence
Time/h
c
No dsRNA control
qa_dsimpa
-QA nuclei
+QA nuclei
Nuclear FRQ
Nuclear FRQ (a.u.)
Hyphal FRQ
Hyphal FRQ (a.u.)
-QA whole cell
+QA whole cell
d
csr-1::qa_dsimpa impα^V5
frq fold changes LP/DD21
wc-1 fold changes LP/DD21
impα fold changes LP/DD21
- QA
+ QA
e
No dsRNA control
qa_dsimpa
-QA nuclei
+QA nuclei
Nuclear WC-2
Nuclear WC-2 (a.u.)
Hyphal WC-2
Hyphal WC-2 (a.u.)
20241019 #1 -QA
20241019 #1 +QA
20241019 #2 -QA
20241019 #2 +QA
20241003 #1 -QA
20241003 #1 +QA
20241003 #2 -QA
20241003 #2 +QA
20241117 #1 -QA
20241117 #1 +QA
20241117 #2 -QA
20241117 #2 +QA

**Supplementary Figure 3 | Deficiency of Impα disrupts the circadian clock by reducing nuclear FRQ and total WC-1.**

a) Full blots of Fig. 3d, with more *qa_dsimpα* strains included. * indicates unspecific bands in immunoblots. b) Luciferase assays of control (*impα*$^{V5}$) or *qa_dsimpα* strains with varying QA concentrations (from Fig. 3e), plotted separately for clearer visualization. c) Quantification of nuclear (top) or total (bottom) FRQ signal in tip regions from 3D renderings of z-stacks, with each biological replicate plotted separately. d) Fold changes of *frq*, *wc-1*, or *impα* after light-pulse with or without $10^{-2}$ M QA, calculated from data presented in Fig. 4e-g. e) Quantification of nuclear (top) or total (bottom) WC-2 signal in tip regions from 3D renderings of z-stacks, with each biological replicate plotted separately.

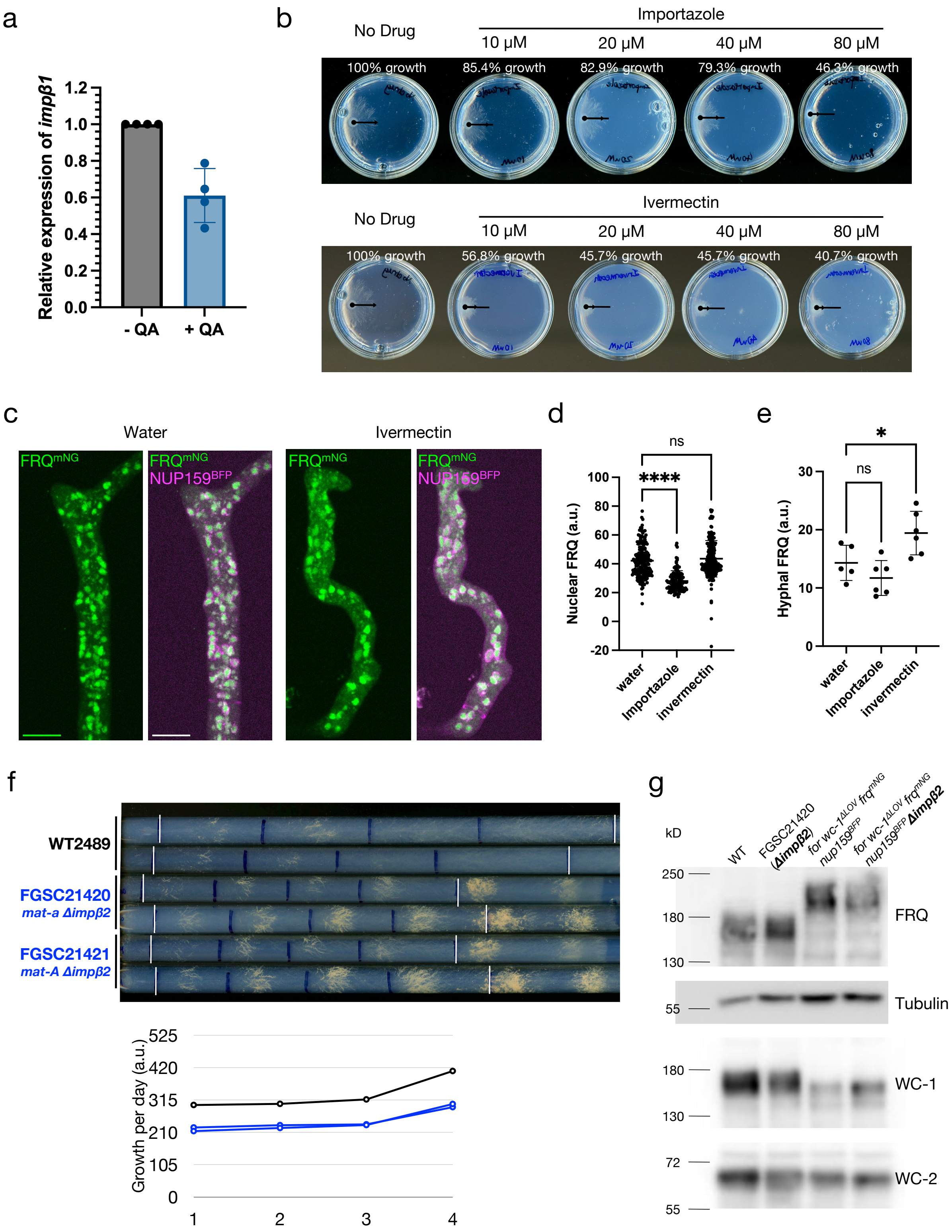

a
Relative expression of impβ1
1.2
1.0
0.8
0.6
0.4
0.2
0.0
- QA
+ QA
b
No Drug
Importazole
10 μM
20 μM
40 μM
80 μM
100% growth
85.4% growth
82.9% growth
79.3% growth
46.3% growth
No Drug
Ivermectin
10 μM
20 μM
40 μM
80 μM
100% growth
56.8% growth
45.7% growth
45.7% growth
40.7% growth
c
Water
Ivermectin
FRQmNG
NUP159BFP
d
Nuclear FRQ (a.u.)
ns
****
water
Importazole
invermectin
e
Hyphal FRQ (a.u.)
*
ns
water
Importazole
invermectin
f
WT2489
FGSC21420
mat-a Δimpβ2
FGSC21421
mat-A Δimpβ2
Growth per day (a.u.)
525
420
315
210
105
0
1
2
3
4
g
kD
WT
FGSC21420 (Δimpβ2)
for wc-1ΔLOV frqmNG nup159BFP
for wc-1ΔLOV frqmNG nup159BFP Δimpβ2
250
180
130
FRQ
55
Tubulin
180
130
WC-1
72
55
WC-2

**Supplementary Figure 4 | Effects of Impβ1 or Impβ2 on circadian clock.**

a) Expression level of *impβ1* in *qa_dsimpβ1* strains with or without $10^{-2}$ M QA quantified by RT-qPCR. Error bar represents the standard deviation from biological triplicates. b) Growth of the wildtype strain (661-4a) after 1 day with varying concentrations of Impβ1 inhibitors. Black dots indicate the inoculation center, black reference lines (all equal length) show growth without drug, and small vertical bars mark the colony growth front. The percentages of growth compared to no-drug condition are calculated. c) Maximum projection images of example z-stacks showing FRQ subcellular localization with or without Ivermectin. d-e) Quantification of d) nuclear FRQ or e) total FRQ signal in tip regions from 3D renderings of z-stacks (mean ± SD) with Ivermectin, compared to Importazole treatment data from Fig. 5b-c. f) Race tube assays of wildtype and *Δimpβ2* strains. The strains were entrained by a light to dark transfer, with white bars marking the beginning and end of 4-day growth under constant darkness, and black bars marking the growth front every 24 hours. The growth of every 24 hours, quantified by distances between bars, is plotted at the bottom. Black circles represent the wildtype strain, and blue circles represent *Δimpβ2* strains. g) Core clock protein levels in *Δimpβ2* strains or their control strains.

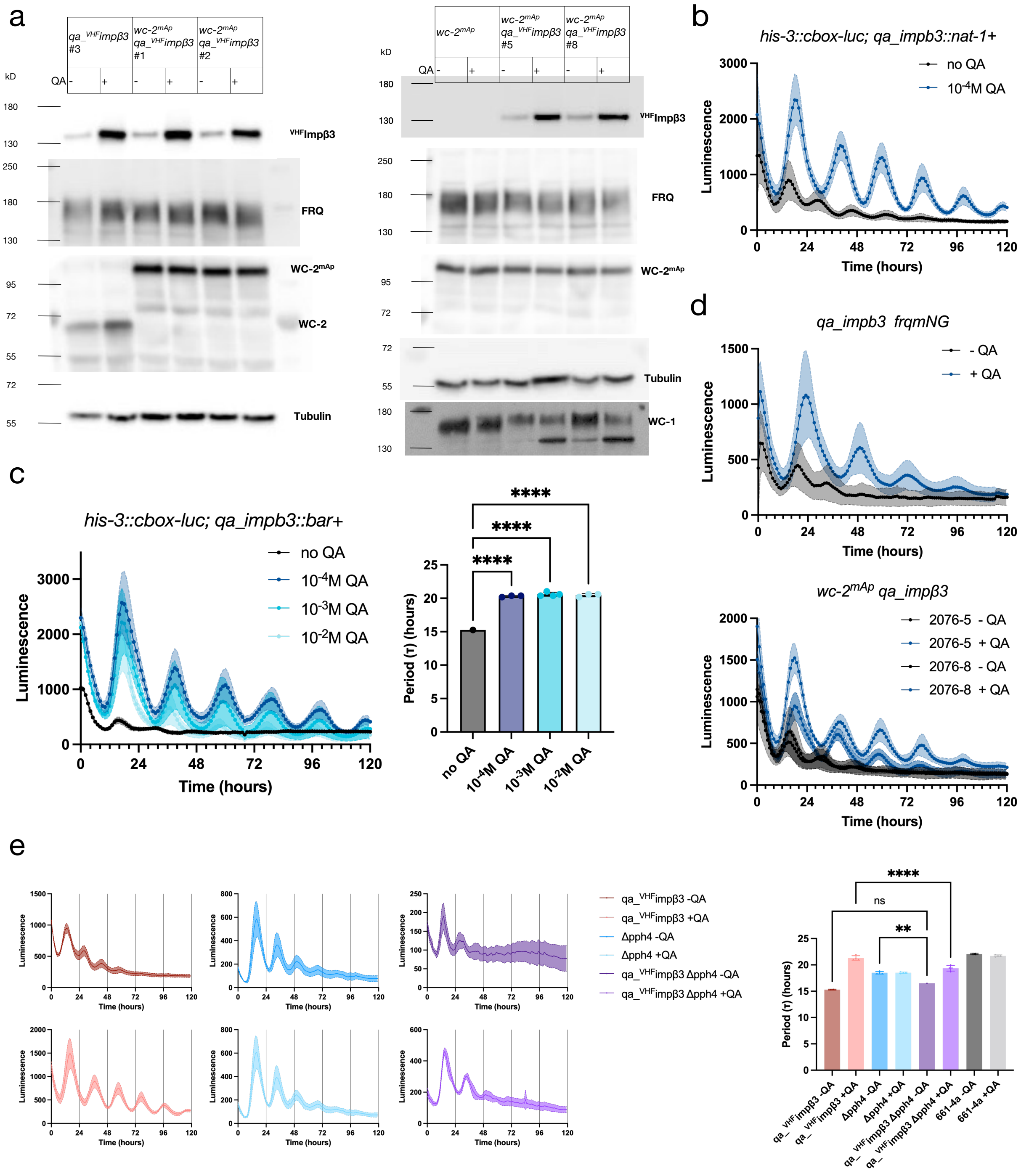

a
qa_VHFimpβ3 #3
wc-2mAp qa_VHFimpβ3 #1
wc-2mAp qa_VHFimpβ3 #2
kD
QA - + - + - +
180
130
250
180
130
95
72
55
72
55
VHFImpβ3
FRQ
WC-2mAp
WC-2
Tubulin
wc-2mAp
wc-2mAp qa_VHFimpβ3 #5
wc-2mAp qa_VHFimpβ3 #8
QA - + - + - +
WC-1
b
his-3::cbox-luc; qa_impb3::nat-1+
no QA
10-4M QA
Luminescence
Time (hours)
c
his-3::cbox-luc; qa_impb3::bar+
no QA
10-4M QA
10-3M QA
10-2M QA
Luminescence
Time (hours)
Period (τ) (hours)
****
no QA
10-4M QA
10-3M QA
10-2M QA
d
qa_impb3 frqmNG
- QA
+ QA
wc-2mAp qa_impβ3
2076-5 - QA
2076-5 + QA
2076-8 - QA
2076-8 + QA
Luminescence
Time (hours)
e
qa_VHFimpβ3 -QA
qa_VHFimpβ3 +QA
Δpph4 -QA
Δpph4 +QA
qa_VHFimpβ3 Δpph4 -QA
qa_VHFimpβ3 Δpph4 +QA
Luminescence
Time (hours)
Period (τ) (hours)
****
ns
**
661-4a -QA
661-4a +QA

**Supplementary Figure 5 | Effects of Impβ3 on circadian clock.**

a) Additional replicates showing core clock protein levels in control strains or inducible *impβ3* strains with or without $10^{-2}$ M QA. b) Luciferase assays of the inducible *impβ3* strain with or without $10^{-4}$ M QA. Each curve represents the average of biological triplicates each with 4 technical replicates, with the shade showing one standard deviation. c) Luciferase assays of another independently constructed inducible *impβ3* strain with varying QA concentrations (left) and their period lengths (right). n=4 replicates. The shade shows one standard deviation. Periods were analyzed by one-way ANOVA with multiple comparisons. d) Luciferase assays of the inducible *impβ3* strains used for FRQ (top) or WC-2 (bottom) imaging with or without $10^{-4}$ M QA. Each curve represents the average of two biological replicates each with 2-3 technical replicates, with the shade showing one standard deviation. e) Luciferase assays of strains with both *pph4* deletion and inducible *impβ3* compared to strains with either mutated allele alone, with or without $10^{-4}$ M QA (left) and their period lengths (right). n=4 replicates. The shade shows one standard deviation. Periods were analyzed by one-way ANOVA with multiple comparisons.
